# Divergent TDP43-regulated and TDP43-independent cryptic splicing in the cortex and spinal cord

**DOI:** 10.1101/2024.03.29.587244

**Authors:** Dwight F. Newton, Rena Yang, Johnny Gutierrez, Jeffrey W. Hofmann, Sreedevi Chalasani, Felix L. Yeh, Anne Biever, Brad A. Friedman

**Affiliations:** Computational Science, Roche Canada, 7070 Mississauga Road, Mississauga, Ontario, Canada, L5N 5M8; Scientific Insights Engineering, Genentech, 1 DNA Way, South San Francisco, CA, USA, 94080; Department of Translational Medicine: OMNI Biomarker Development, Genentech, 1 DNA Way, South San Francisco, CA, USA, 94080; Department of Research Pathology, Genentech, 1 DNA Way, South San Francisco, CA, USA, 94080; Department of OMNI Bioinformatics, Genentech, 1 DNA Way, South San Francisco, CA, USA, 94080

**Author notes:** Corresponding author Brad A. Friedman. Contributions BAF and DFN conceptualized the study. DFN and RY performed bioinformatics analyses. JWH selected the spinal cord cohort samples, and analyzed the microscopy data. JG performed molecular biology experiments (tissue processing, RNA extraction, RT-qPCR). SC performed Basescope ISH+IHC experiments. The first draft of the manuscript was written by DFN and all authors commented on subsequent versions of the manuscript. BAF provided oversight to the study including experimental design, data interpretation, and manuscript preparation. All authors read and approved the final manuscript. Declaration of interests All authors are employees of either Genentech Inc. or Hoffman-LaRoche Ltd.

**Keywords:** TAR DNA-binding protein 43 (TDP-43), Amyotrophic Lateral Sclerosis (ALS), Frontotemporal Dementia (FTD), Cryptic splicing, RNA-sequencing

## Abstract

Mislocalization of the nuclear TAR DNA-binding protein 43 (TDP43) is a hallmark of ALS and FTD which leads to de-repression and inclusion of cryptic exons (CEs), promising biomarkers of TDP43 pathology in a spectrum of neurodegenerative diseases. However, most CEs to date have been identified from *in vitro* models or a single cortical FTD dataset, and little is known about cryptic splicing in the spinal cord, or within different neuronal subtypes. We meta-analyzed published bulk RNAseq datasets representing 1,778 RNAseq profiles of ALS and FTD post-mortem tissue, and *in vitro* models with experimentally depleted TDP43. We identified 142 cryptic splices, including 68 novel events. We found divergent cryptic splicing primarily between the spinal cord and cortex, validated in an independent ALS cohort by qPCR and supported by *in situ* hybridization (ISH). We also identified a set of cryptic splices observed in tissue but not *in vitro*, and, being present in either SOD1-ALS or MAPT-FTD subjects, likely TDP43 independent. Finally, leveraging multiple public single-nucleus RNAseq datasets of ALS and FTD motor and frontal cortex, we confirmed the elevation of cortical-enriched splices in disease and localized them to layer-specific neuronal populations. We provide a web interface to browse the meta-analysis results at https://go.roche.com/CrypticSplicingLandscape. This catalog of cryptic splices will inform efforts to develop biomarkers for tissue-specific and cell type-specific TDP43 pathology.

## Introduction

Abnormal processing of the protein TDP43 is observed in post-mortem tissue from a spectrum of neurodegenerative diseases, including amyotrophic lateral sclerosis (ALS) and frontotemporal dementia (FTD)[1]. Although normally localized to nuclei, in pathological contexts this splicing factor accumulates in primarily cytoplasmic aggregates. To understand the molecular consequences of nuclear TDP43 loss-of-function, several groups have performed RNAseq profiling on *in vitro* models with experimentally depleted TDP43 levels [2–5]. These studies identified the accumulation of mRNA transcripts including so-called “cryptic” splice junctions not normally included in mature transcripts [6,7]. Some of these cryptic splices, namely in the genes STMN2 and UNC13A, were subsequently observed in RNAseq profiling of FTD and ALS post-mortem tissue, but not healthy controls [2–5], and were enriched in TDP43-deficient neuronal nuclei sorted from ALS/FTD frontal cortex [8].

This represents powerful evidence demonstrating that the loss of TDP43 splicing repression results in the accumulation of cryptic splices and is a defining pathological hallmark of these proteinopathies.

However, clinical biomarkers that can directly diagnose TDP43 pathology in nervous system tissues from accessible patient biosamples do not yet exist, though reports of ALS/FTD extracellular vesicles suggest potential utility[9]. Thus, regardless of whether cryptic-spliced mRNAs contribute to the pathogenesis of ALS or FTD, or are merely consequences of a primary proteinopathy, they are potentially of high value as biomarkers. Given their virtual absence in healthy conditions, the detection of even low levels of cryptic mRNA or translated protein in biofluids could facilitate diagnosis and disease monitoring in patients. Furthermore, such biomarkers could enable accurate screening for patient inclusion (or exclusion), or pharmacodynamic evaluation for therapeutics targeting TDP43 pathology. Pharmacodynamic biomarkers could be particularly impactful to enable validation of TDP43-modifying agents in smaller cohorts preceding large, expensive registrational trials

An ideal biomarker assay would measure, likely via RNA or encoded peptide, the levels of one or a small number of all of the possible cryptic splices. Indeed, recent studies have shown the presence of translated cryptic peptides in ALS cerebrospinal fluid[10], and the elevation above healthy controls of a cryptic peptide in the gene HDGFL2 in CSF of some C9orf72+ ALS subjects[11]. However, it is not clear that all cryptic splices are equally sensitive to TDP43 pathology, detectable, or have similar effects on gene or protein expression. Some may be very sensitive to loss of TDP43 activity, whereas others might require deeper TDP43 repression. Some cryptic splices may be located in non-coding exons, or result in unstable proteins, precluding peptide-level detection. Finally, even for an RNA or protein which stably accumulates in tissue, detection in accessible biosamples such as plasma or cerebrospinal fluid depends on transport to, and stability in, these compartments; properties which are very hard to predict. Therefore, having a complete catalog of cryptic splices will be essential to identify the optimal targets to establish robust assays.

Aside from purely physiological challenges, the potential universe of cryptic splices could depend on the cellular context, due to differences in expression of cryptic splice-harboring genes or the activity of splicing factors other than TDP43. Therefore, certain cryptic splices might be specific to spinal cord motor neurons lost in ALS, while others might reflect pathological processes in cortical neurons or other cell types, making them potentially useful in monitoring different subtypes of TDP43 pathology. Identifying context-specific cryptic splices has represented a particular challenge as most sequencing-based screens have focused on *in vitro* models exclusively [2,4,5] or a limited set of *in vivo* data (primarily from ALS/FTD frontal cortex) [8]. These approaches, while allowing for a substantial enrichment of signal, may not be generalizable to the wider array of nervous system tissues affected by TDP43 pathology.

We therefore set out to create an expanded catalog of cryptic splices by meta-analyzing published RNAseq datasets from *in vitro* neuron models and post-mortem brain and spinal cord tissue from ALS and FTLD-TDP subjects, comprising 1,778 bulk RNAseq samples across 16 contexts. Our cryptic splice discovery pipeline [12,13] identified 142 events across these datasets, including 68 which were not reported in previous analyses. Comparison of these cryptic splices across datasets revealed significant differences between cryptic splicing in cortical tissue, spinal cord, and *in vitro*. We validated a subset of these novel events with multiplex RT-qPCR on RNA extracted from spinal cord and cortical samples of controls and ALS subjects with histopathologically confirmed TDP43 pathology, and spatially validated selected splices with *in situ* hybridization. Using several published single-nucleus RNAseq datasets of ALS and FTD cortex, we confirmed the elevation of cortical-enriched cryptic splices in disease, and localized them specifically to L2/3, L5, and L6 excitatory neurons.

## Results

### Post-mortem and in vitro transcriptomics datasets reveal diverse splicing patterns

To identify cryptic splices associated with ALS, frontotemporal lobar degeneration with TDP43 pathology (FTLD-TDP), and experimental TDP43 depletion (TDP43-KD), we leveraged publicly available bulk RNAseq datasets (Table 1) comprising 1,871 samples (1,778 after QC filtering) of post-mortem patient RNA or sorted nuclei across 14 CNS regions and *in vitro* cell lines (Figure 1a) [4,5,8,14–17]. The post-mortem RNA samples included cerebellum, hippocampus, cortex and spinal cord from ALS, FTLD-TDP, FTLD-Tau, and control subjects generated by the New York Genome Center (NYGC) ALS consortium[4,14,15,17] and, separately, the RiMOD-FTD consortium [16]. We also included a study comparing TDP43+ and TDP43- neuronal nuclei from ALS/FTLD-TDP frontal cortex (hereafter referred to as “Liu”) [8]. Finally, four datasets using siRNA-based TDP43-KD in different models were included [4,5].

**Figure 1.**
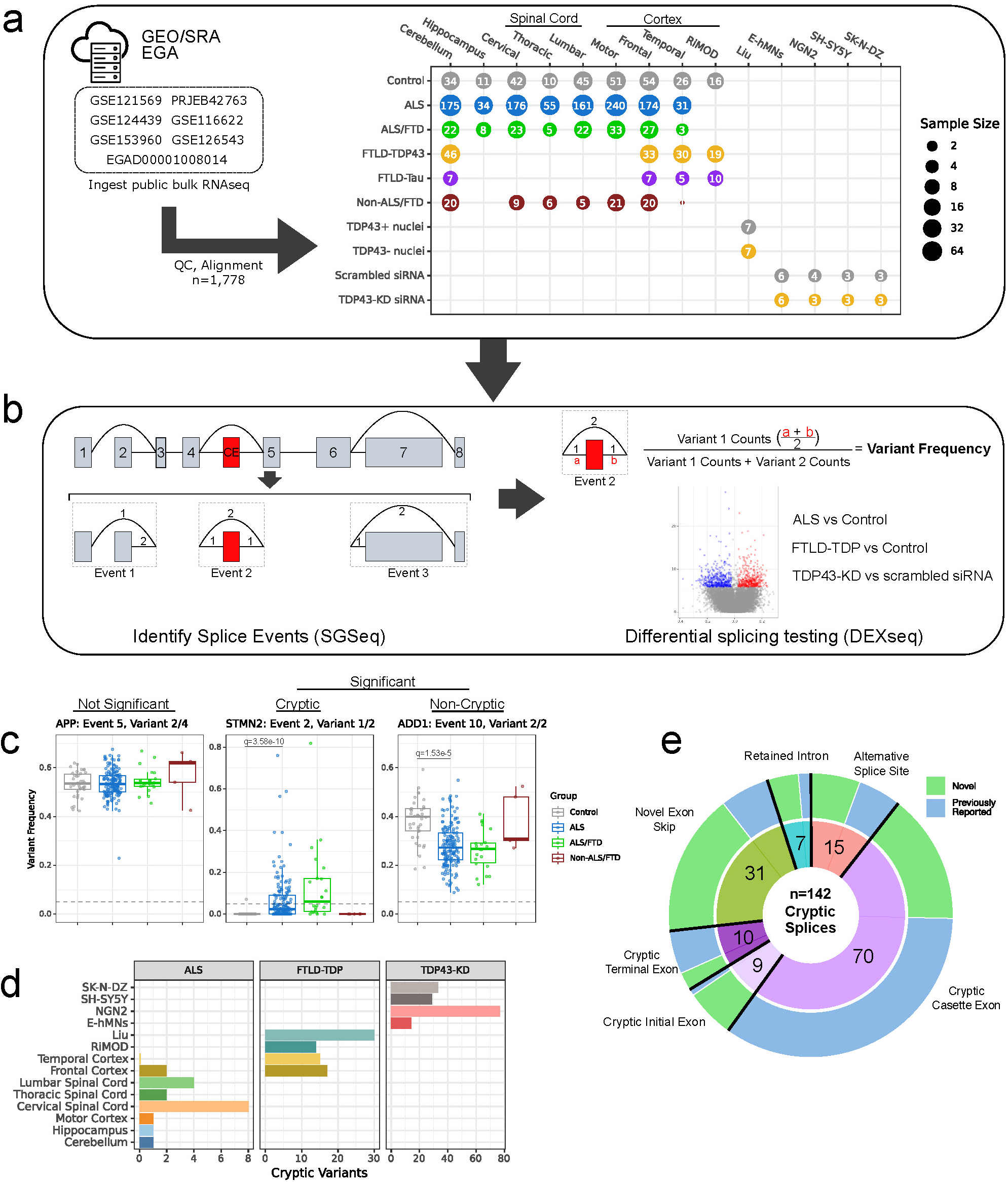
**Analytical workflow used to meta-analyze cryptic splicing in post-mortem and *in vitro* RNAseq**. **a** - Data sources and respective sample sizes by experimental group. Named tissues (Cerebellum through Temporal) are all from the NYGC ALS Consortium; RiMOD consortium data are from the frontal cortex. **b** - Conceptual framework used in SGSeq to identify splicing events (left) and test for differential splicing using DEXseq (right). Known transcript models and novel exonic and splice junction features identified from input RNAseq data are used to build a splice graph describing the potential splicing of a given gene. Splice events are identified as regions of the splice graph with multiple potential paths sharing start and/or end nodes. Three example events are shown representing (1) an alternative first exon, (2) a cassette exon (with a CE inclusion), and (3) an alternative last exon. Within a splice event, each path is termed a splice variant (here, identified using small numbers annotating features), and variant frequencies for each are calculated. DEXseq is then used to identify splice variants with significant differences in variant reads across contrasts of interest. **c** - Representative examples of our categorization of splicing. Each point shows the variant frequencies in a single lumbar spinal cord sample. Grey dashed lines indicate control variant frequency threshold of 0.05. Events and variant numbers correspond, broadly, to 5’ to 3’ order along gene length. **d** - Number of cryptic splice variants identified in each differential contrast performed in context-specific analyses. CNS region/dataset/model is indicated on the y-axis, and differential contrast indicated on subplot titles. **e** - Overview of cryptic splices identified in this study, across all contrasts, categorized by splice type. Splices categorized as previously-reported have been described in one or more of Brown et al (2022), Ma et al (2022), Klim et al (2021), Seddighi et al (2023), Humphrey et al (2017), or Ling et al (2015).

We re-analyzed these datasets from raw FASTQs, utilizing a single analytical framework employing the *GSNAP* algorithm [18,19] for read alignment followed by *SGSeq* and *DEXseq* R packages for splice detection and differential testing, respectively [12,13]. We organized the 7 datasets into 16 “contexts” representing different tissue regions, data sources (e.g. NYGC and RiMOD-FTD frontal cortex were separate contexts), diseases, or different *in vitro* models, analyzing each context independently. Briefly, the SGSeq framework generated a splice graph using ENSEMBL-annotated GRCh38 transcripts supplemented with novel exonic and intronic features observed from RNAseq alignments (Figure 1b). Variable splice graph regions with shared start and end nodes (*splice events*) were identified (Figure 1b) and the relative usage of paths through each event (*splice variants*) was quantified as the *variant frequency* (VF) for each sample, a number between 0 and 1 (equivalent to *percent spliced in* - Ψ, which ranges from 0-100%). Across the 16 differential contrasts performed (see Methods), we calculated the difference in variant frequency (ΔVF) between controls and ALS, FTLD-TDP, TDP43-KD, or TDP43- nuclei, as appropriate. We set differential splicing significance at |ΔVF| > 0.025, and 5% false discovery rate (FDR).

Similar to previous studies[2,10], we filtered these differentially spliced variants to keep only those with minimal inclusion in controls (defined here as mean or median VF < 0.05 in controls, Figure 1c). Thus, in this meta-analysis, cryptic splices were defined by the statistical framework of being significantly differentially spliced in disease and showing very low, or no, inclusion in controls.

The number of cryptic splices observed in each dataset broadly agreed with regional expectations of TDP43 pathology: FTLD-TDP frontal and temporal cortex greater than ALS, ALS motor cortex greater than FTLD-TDP, and ALS spinal cord greater than non-spinal cord (Figure 1d). Cryptic splice events (shortened hereafter to “cryptic splices”) were consolidated across contexts, yielding a catalog of 142 cryptic splices, consisting largely of CE inclusion events (70 cassette, 10 terminal, and 9 novel start exons), 31 exon skipping events, and a small number of retained introns and alternative splice sites (Figure 1e), with distinct tissue, disease, and model-specificity. 133/142 cryptic splices contained unannotated splice junctions. Our GSNAP/SGSeq analysis gave broadly concordant results to the STAR/MAJIQ-based approach previously published for NGN2-neurons [4] (Online Resource 1).

### Cryptic splicing exhibits spinal, cortical, and in vitro-enriched signatures

To understand if the 142 identified cryptic splices showed context specificity across the included datasets we performed hierarchical clustering, which revealed six distinct clusters representing disease, tissue, and model-specificity (Figure 2a, Online Resource 2). Distinct from these six splice sets, inclusion of the STMN2 CE 2a was unique in being elevated in all contexts in our RNAseq meta-analysis: ALS spinal cord, FTLD-TDP and ALS cortex, sorted TDP43- neurons, and all TDP43-KD models. The SGSeq-predicted cryptic junction coordinates of STMN2 aligned precisely with those reported previously (Figure 2b - left) [3]. We identified a spinal cord-enriched cryptic splice signature (7 splice variants), which was virtually absent from other CNS regions with TDP43 pathology or *in vitro* TDP43-KD models. One such splice was identified in CHCHD6, representing potential usage of an alternative first exon (Figure 2b - center). An *in vivo*-enriched signature (6 splice variants) was elevated across multiple post-mortem contexts (across both FTLD-TDP43 and FTLD-Tau subjects) but not in *in vitro* models. A cortical splice signature (Cortical1 - 24 splice variants) was elevated in ALS-FTD/FTLD-TDP frontal/temporal cortex, ALS motor cortex, and *in vitro* models. This cortical signature included a novel terminal CE in HS6ST3 (Figure 2b - right), as well as the well-documented cryptic cassette exon in UNC13A (Online Resource 3)[2]. An *in vitro-*enriched signature (27 splice variants) was particularly evident in neuroblastoma cell lines, and largely absent *in vivo*. 39 splice variants formed a signature, showing enrichment in FTLD-TDP and *in vitro* models, but not post-mortem ALS, which we named “Cortical2”. Lastly, a neuronal signature of 38 splice variants was enriched across TDP43-KD models and, partly, in TDP43-negative neuronal nuclei from FTLD-TDP frontal cortex. Examples of splices from each signature are highlighted in Online Resource 3.

**Figure 2.**
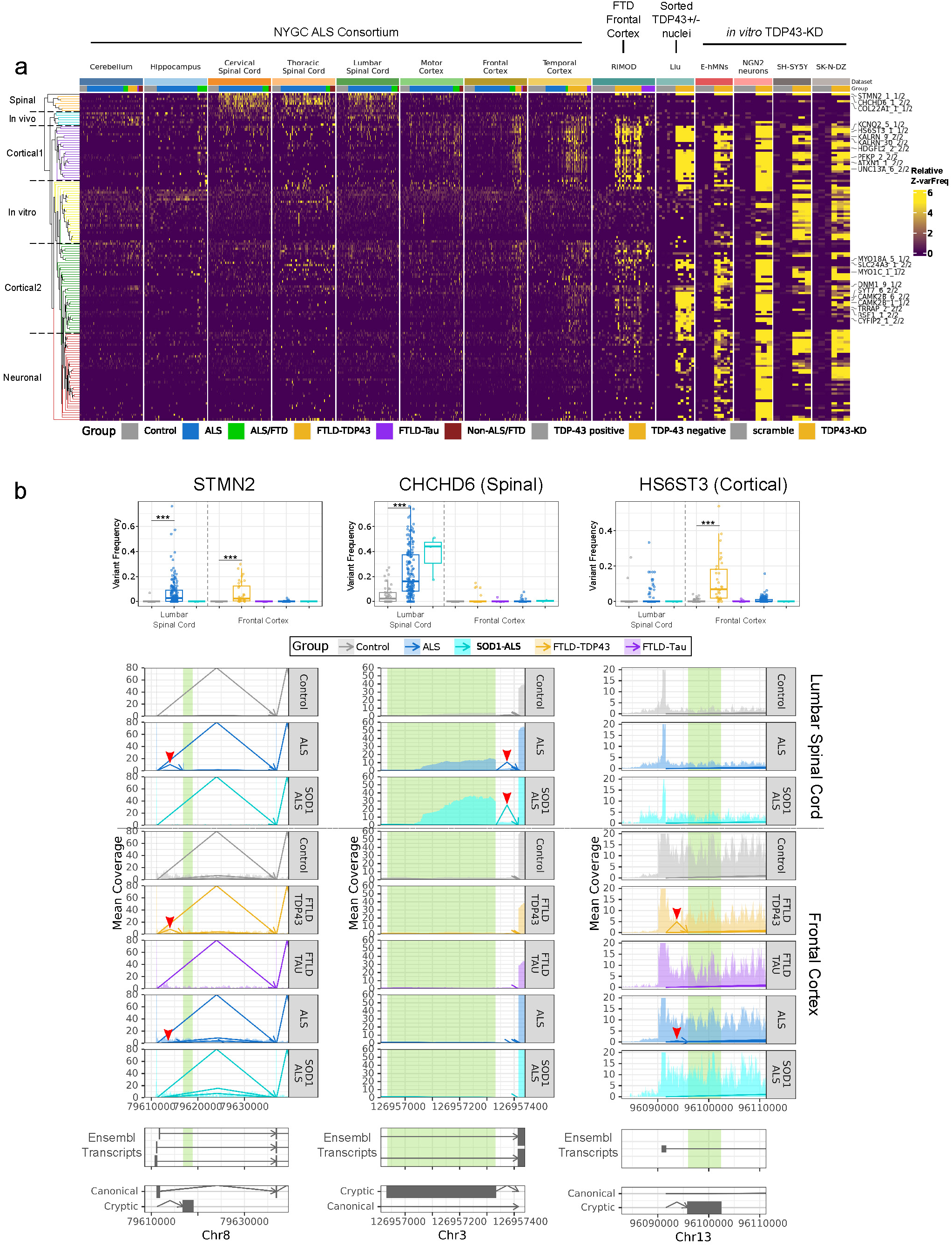
Cryptic splicing shows context-specificity, particularly between cortex and spinal cord. **a** - Heatmap showing inclusion of 142 cryptic splices across all datasets included in this study. Variant frequency was Z-normalized (see methods) to the mean and standard deviation of controls of each context for visualization purposes. Control groups are colored in grey, and contexts with TDP43 depletion confirmed by experimental condition, *post mortem* histopathology, or presence of genetic factors are colored in gold. Source data:Online Resource 20. **b** - Variant frequency and coverage plots of three illustrative cryptic splices for different splice sets. Variant frequency plots (above legend) contain the sample-wise quantification of the cryptic variant denoted in the coverage plots.*** indicates a DEXseq FDR-corrected q-value < 10^-6^. Next, “Mean Coverage” track shows mean RNAseq coverage, aggregated by group and tissue, in histograms and splice junctions with arrows. Junction coordinates are indicated by arrow start and end positions, with the height indicating the junction read depth, and strand indicated by position of arrow head (minus strand left, plus strand right). Key cryptic junctions are highlighted with red arrow heads. For visual clarity, junctions with less than 10 reads across all samples, and those on the opposite strand as the gene of interest, are not shown. CE coordinates are highlighted in green. “Ensembl Transcripts” track shows canonical Ensembl transcripts present in the same coordinate space (rectangles: exons, arrows: introns, with arrows pointing from splice donor to acceptor site). Bottom track shows the cryptic and canonical splice events identified by SGSeq, in which arrows indicate splice junctions and boxes indicate exonic features. Note the spinal cord-specificity of the CHCHD6 cryptic splice (with inclusion in SOD1-ALS), cortical-specificity of the HS6ST3 cryptic splice, and the presence of the STMN2 cryptic splice in both regions.

As our definition of “cryptic” (mean or median VF < 0.05 in controls) is context-specific, we considered whether any cryptic events might be non-cryptic (VF ≥ 0.05 in controls) in another context. We found that 16/142 splices exhibited broad inclusion across *in vivo* datasets (mean VF of cryptic splice > 0.1) regardless of diagnostic group (Online Resource 4). Critically, all such splices were discovered from *in vitro* analyses, and 11/16 were members of *in vitro* splice sets, with 5 in the Cortical2 splice set (which is also enriched in *in vitro* models). We omitted these splices in downstream analyses, such as the generation of splice scores, but retain them in our catalogue given they reflect cryptic splicing within a particular *in vitro* context. No cryptic splices derived from *in vivo* analyses were non-cryptic *in vitro*.

Readers can find in-depth reports for each cryptic splice, showing the inclusion across all meta-analysis studies, variant structure and coverage plots, precise genomic coordinates, inclusion in GTEx v6, and snRNAseq studies of ALS/FTD at our companion website “The Cryptic Splicing Landscape” (https://go.roche.com/CrypticSplicingLandscape). All cryptic splicing quantifications are provided inOnline Resource 20.

### Spinal splice set does not show evidence of TDP43-regulation, but is ALS-specific

Using whole-genome sequencing information provided by the NYGC, we identified a small group (n=3) of ALS donors which harbour mutations in SOD1. This small population, which does not harbour TDP43-pathology[20], showed similar levels of cryptic CHDHD6 inclusion to non-SOD1 ALS (Figure 2B, centre), as did all other splices in the spinal splice set (Online Resource 5), suggesting these are not directly TDP43-regulated. Inclusion of the *in vivo* cryptic splices, which also showed inclusion in the spinal cord, was largely not elevated in SOD1-ALS (Online Resource 5). Expanding beyond the NYGC spinal cord cohort, the spinal splice set was elevated in ALS subjects in two other studies performing RNAseq profiling cervical spinal cord and lumbar anterior horns (Online Resource 6).

Positing this profile may be reflective of a general, pan-disease, neurodegenerative processes, we examined inclusion of the spinal splice set in RNAseq profiling of spinal cord injury and disease (intervertebral disc degeneration, spinal muscular atrophy), and other neurodegenerative diseases including MS and AD [21–26]. Cryptic splicing analysis of these datasets showed virtually no broad inclusion of the spinal cryptic splices, highlighting their ALS-specificity (Online Resource 7a). Quantification of the spinal cryptic splices in the Genome-Tissue Expression (GTEx) project[27], a vast database of >6,000 healthy RNAseq profiles across the human body, found spinal splices included at low-to-moderate levels in the skin, lung, and immune cell-rich tissues (spleen and blood), depending on the splice in question (Online Resource 7b). COL22A1 and MYO5A cryptic splices showed virtually zero inclusion in GTEx, similar to other *bona fide* cryptic splices such as STMN2 and KALRN (Online Resource 7b).

Lastly, we integrated our spinal cord RNAseq with a recent report from the NYGC characterizing glial/inflammatory and oxidative stress-related molecular subtypes of ALS [28]. 92% (262/283) of our lumbar spinal cord samples overlapped with this study, and we found that all spinal cryptic splices were significantly elevated in the ALS-Glia phenotype versus controls and 5/7 were significantly elevated compared to the ALS-Ox phenotype (Online Resource 7c). Taken together, these results highlight the ALS-relevance of this splicing signature, despite no evidence of TDP43 regulation, and suggest that it may be reflective of an ALS-specific neuroinflammatory state.

### Context-specific gene expression may explain cortical cryptic splicing, but not other signatures

We next explored potential mechanisms for context-specific splicing. One potential explanation could be context-specific expression of the genes which harbor cryptic splices (“host genes”). For each cryptic splicing signature, we calculated the difference in host-gene expression (Figure 3a) in control conditions, across the same datasets showing splicing enrichment. We then computed the percentile of the fold-changes relative to the entire genome to determine the relevance of these differences. No host gene was unexpressed outside the expected context, though 25/136 genes were above the 95th percentile of genome-wide LFCs (Figure 3b). These genes were largely (18/25) from the cortical1 and cortical2 splice sets, indicating that greater expression of these genes in the cortex could explain some of their context-specific splicing. The remaining signatures showed either minimal, or even inverse (spinal), context-enrichment of host genes. Using snRNAseq of healthy spinal cord and cortex from the Human Brain Cell Atlas project (an atlas of 26.3M cells)[29], we observed similar cortical1 host gene enrichment in cortical brain regions, driven by neuron populations (Online Resource 8). Thus, differential host gene expression across contexts explains some cortical-enriched cryptic splicing, but not the majority of context-specific cryptic splicing.

**Figure 3.**
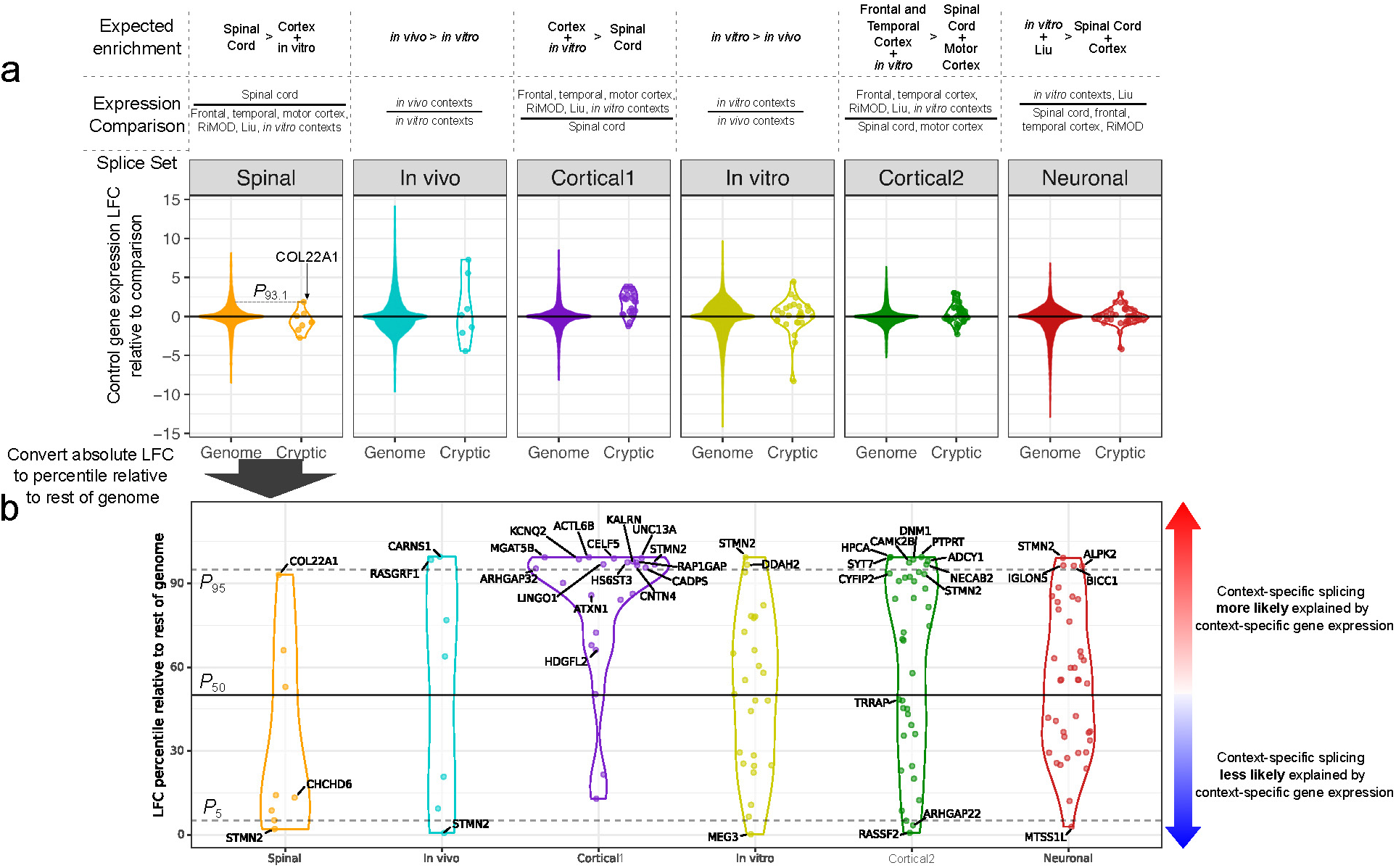
Context-specific cryptic splicing is modestly related to context-specific host gene expression. Cross-context cryptic splice host gene expression differences compared to genome-wide distributions. **Spinal splice set**: cervical, thoracic, and lumbar spinal cord versus frontal, temporal, motor cortex, RiMOD, Liu (sorted TDP43+ nuclei), and all *in vitro* scrambled controls. ***In vivo* splice set**: all spinal cord and cortical regions (including RiMOD, but not Liu) versus all *in vitro* controls. **Cortical1 splice set**: frontal, temporal, motor cortex, RiMOD, Liu, and all *in vitro* controls vs all spinal cord controls. ***In vitro* splice set**: all *in vitro* controls versus all spinal cord and cortical regions. **Cortical2 splice set**: frontal, temporal cortex, RiMOD, Liu, and all *in vitro* controls versus spinal cord and motor cortex controls. **Neuronal splice set**: Liu + all *in vitro* controls versus all spinal cord and cortical regions. **a**: Log_2_ fold-change (LFC) in expression of cryptic splice host genes (“cryptic”) within each splice set and all other genes (“genome”), in control subjects of observed contexts of enrichment compared to controls of non-enriched contexts. Dashed line indicates how LFC percentiles were determined using spinal splice COL22A1 as an example. The dashed line indicates that the inter-context LFC of COL22A1 (1.86) corresponded to the 93.1 percentile of all inter-context LFCs. **b**: LFC percentile for each cryptic splice, relative to the genome-wide LFC distribution for each comparison of contexts. Solid horizontal line indicates 50th percentile and dashed lines indicate 5th and 95th percentiles. Genes with LFCs beyond the 5th and 95th percentiles, and those significantly elevated in qPCR experiments (including STMN2 for reference), were labeled.

Another source of context specificity, particularly for the spinal splice set, may be tissue-specific enrichment of trans-regulatory factors such as RNA binding proteins (RBPs) beyond TDP43. We collected many published cross-linking and immunoprecipitation sequencing (CLIPseq) repositories, representing 570 CLIPseq peaksets across 259 RBPs [30–32]. Localizing these CLIPseq peaks to the canonical TDP43 binding region (500bp up-or down-stream of a TDP43-regulated splice site)[33], revealed that the majority of cryptic splices (98/142) showed evidence of a TDP43 CLIPseq peak, or GT-rich sequence (Online Resource 9). Of the spinal splices, only CHCHD6 showed a TDP43 CLIPseq peak in this region. Searching across all included RBPs, CSTF2T showed peaks proximal to the STMN2, ADCY3, IQSEC1, C20orf194, and AFF3 splices, which was enriched above the rate observed in other splices by hypergeometric test (4/6 spinal vs 36/134 non-spinal, p = 0.0025) (Online Resource 10a). However, CSTF2T gene expression did not show spinal enrichment or depletion, and it is thought to regulate polyadenylation site selection [34], not splicing, making it an unlikely regulator of the spinal splicing signature (Online Resource 10b).

### Cryptic splicing has diverse impacts on gene expression and predicted transcript processing

We next explored the relationships of cryptic splicing to host gene expression and transcript stability, given literature observations of cryptic splicing resulting in reduced expression (e.g. STMN2, UNC13A)[2,3] and premature termination codon (PTC) inclusion leading to nonsense-mediated decay (NMD)[35]. We performed context-wise correlations of cryptic splice VF with log-normalized expression, considering relationships significant at 5% FDR (Online Resource 11a). Most (224/230) significant correlations were negative, with only 6 positive VF-expression correlations. However, though most *significant* associations were negative, most correlations were not significant (1016/1246), even when only considering instances where cryptic splicing was transcriptome-wide significant by DEXseq (Online Resource 11a-b). Moreover, splices predicted to result in NMD-sensitive transcripts did not show a greater rate of negative VF-expression correlations (Online Resource 11c). While splices such as STMN2 showed VF-expression anti-correlations in all cases where the splice was significantly elevated, other splices (e.g. KALRN) showed no evidence (Online Resource 11d). Thus, when cryptic splicing is associated with host gene expression it is almost always negative, however most cryptic splices are not in fact significantly correlated with their host gene expression.

### Validation of context-specific cryptic splicing by RT-qPCR, RNAseq, and ISH

To confirm our bioinformatically-predicted cryptic splices, we developed RT-qPCR assays targeting the cryptic variants of 67 cryptic splice events (Online Resource 22) showing *in vivo* inclusion, and quantified their expression in an independent cohort of post mortem lumbar spinal cord and motor cortex from ALS and control subjects (Figure 4a, Table 2). pTDP43 S409/410 immunohistochemistry (IHC) was performed in neighbouring tissue sections to those used for qPCR. pTDP43 staining was classified into 4 categories ranging from 0 (no pTDP3 aggregates) to 3 (many cells showing abundant pTDP43 aggregates), by a neuropathologist blinded to the donor diagnoses (see methods, Figure 4a). 26 cryptic splices were significantly elevated at 5% FDR in ALS, with 14 elevated in both spinal cord and motor cortex, 7 being spinal-specific (including 4 spinal splices), and 5 cortical1 splices being motor cortex-specific (Figure 4b). No cryptic splices were elevated in ALS/FTD motor cortex (which had low pTDP43 pathology) except for HS6ST3. Correlations between cryptic splice qPCR expression and pTDP43 staining further supported this combination of context-specific and context-independent cryptic splicing. The spinal splices correlated with pTDP43 in the spinal cord, the cortical1 splices showed like enrichment in the motor cortex, and the cortical2 splice set showed a cluster of splices correlating with pTDP43 in each region (a small subset of the cortical1 and spinal splices were pTDP43-associated in the opposing region) (Figure 4c). We also performed RNAseq on the spinal cord samples, and found STMN2 and CHCHD6 cryptic splices were significantly elevated in ALS samples, and correlated with both pTDP43 staining and qPCR-based expression (Online Resource 12).

**Figure 4:**
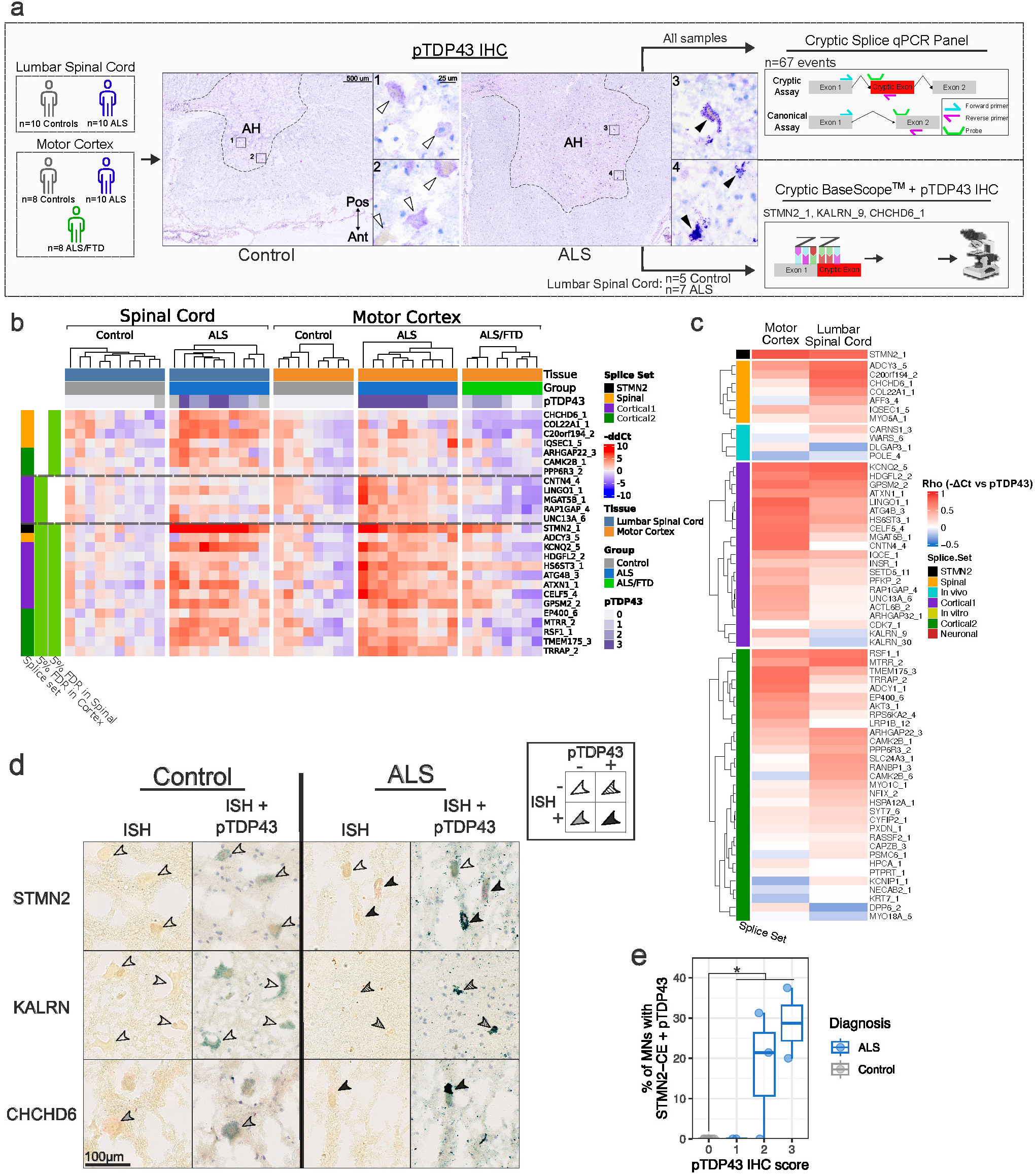
Validation of spinal cord and cortex-specific cryptic splicing in an independent ALS cohort. **a** - Validation strategy and experimental workflow. ALS and control lumbar spinal cord and motor cortex cohorts were assessed for TDP43 pathology by pTDP43 S409/410 IHC. Representative images of pTDP43 IHC in a control (left panel) and ALS (right panel) lumbar spinal cord section. Anatomical landmarks (AH - anterior horn; dashed line - grey/white interface; all images oriented with anterior side at the bottom) are noted on low-magnification images. Boxed regions of interest are highlighted in high-magnification images (right of each panel), showing strong non-nuclear intense staining of pTDP43 in ALS motor neurons (black arrows), and only diffuse background stain in control motor neurons (white arrows). Scale bars: 500 um (large images), 25 um (small images). RNA was extracted from neighbouring sections and RT-qPCR against 67 cryptic splice variants was performed. A subset of spinal cord samples with confirmed TDP43 pathology were used for spatial validation by sequential cryptic splice junction-specific ISH followed by pTDP43 IHC. Some image assets created in BioRender. Newton, D. (2025) https://BioRender.com/18wh7kv. **b** - Heatmap of-ΔΔCt values calculated relative to the mean of HPRT1 and GAPDH loading controls, and then normalized to the mean of control samples (see methods). Only cryptic splices significantly elevated (unpaired Wilcoxon test, 5% FDR) in either spinal cord or motor cortex cohorts are shown. **c** - Heatmap of spearman correlations between cryptic splice expression by qPCR and pTDP43 staining. **d** - Representative images from cryptic ISH+IHC experiments, showing motor neurons in the anterior horn. Arrowheads indicate examples of pTDP43 and/or cryptic-ISH positive or negative cells. **e** - Quantification of STMN2 cryptic ISH signal. Y-axis shows the percentage of all motor neurons in the anterior horn that are positive for both pTPD43 and STMN2-cISH. Some assets in **A** created in BioRender. Newton, D. (2025) https://BioRender.com/18wh7kv

To spatially localize selected cryptic splices in ALS and control tissues, we designed Basescope™ probes targeting the cryptic splice junctions of STMN2 (context-independent), KALRN_9 (cortex-enriched), and CHCHD6 (spinal-enriched). Chromogenic ISH followed by pTDP43 staining on the same slide allowed co-localization of these stains after image alignment. STMN2 cryptic ISH (STMN2-cISH) co-localized with pTDP43 staining in ALS subjects but not controls (Figure 4d - top row), with significantly greater pTDP43+STMN2-cISH positive cells detected in ALS, versus zero in controls (Figure 4e). KALRN-cISH was fully undetectable in either control or ALS spinal cord, consistent with RNAseq and qPCR findings (Figure 4d - second row). CHCHD6-cISH was observed in both control motor neurons, and pTD43+ neurons in ALS (Figure 4c - third row), suggesting a potential non-neuronal origin for the elevation observed by qPCR. Taken together, these qPCR results are consistent with our RNAseq-based description of context-specific cryptic splicing, highlighting previously unseen non-TDP43-regulated cryptic splices in the spinal cord with a more sensitive method of detection. We also identify additional cross-context cryptic splices beyond STMN2 in ADCY3, KCNQ2, HDGFL2, HS6ST3, ATG4B, ATXN1, CELF5, GPSM2, EP400, MTRR, RSF1, TMEM175 and TRRAP.

### snRNAseq localizes cryptic splicing in cortex to L2/3, L5, and L6 IT excitatory neurons

Finally, we sought to determine the cell-type enrichment of cryptic splices, utilizing snRNAseq datasets from two recent studies profiling (1) motor cortex from sporadic and C9orf72 ALS and FTLD patients (Pineda et al.) [36] and (2) motor and frontal cortex (analyzed separately) from C9orf72 ALS and FTLD patients (Li et al.) [37]. One previous study detected STMN2 and KALRN cryptic splices by snRNAseq[38], and we have expanded our search here to include our catalogue of cryptic splices and more datasets. After QC and mutual-nearest-neighbor batch correction [39], 415,741 nuclei were retained from Pineda yielding 29 clusters (Figure 5a). Likewise, analysis of Li motor cortex yielded 59,610 cells forming 31 clusters (Online Resource 15a), while the frontal cortex dataset comprised 63,076 nuclei forming 34 clusters (Online Resource 19a).

**Figure 5.**
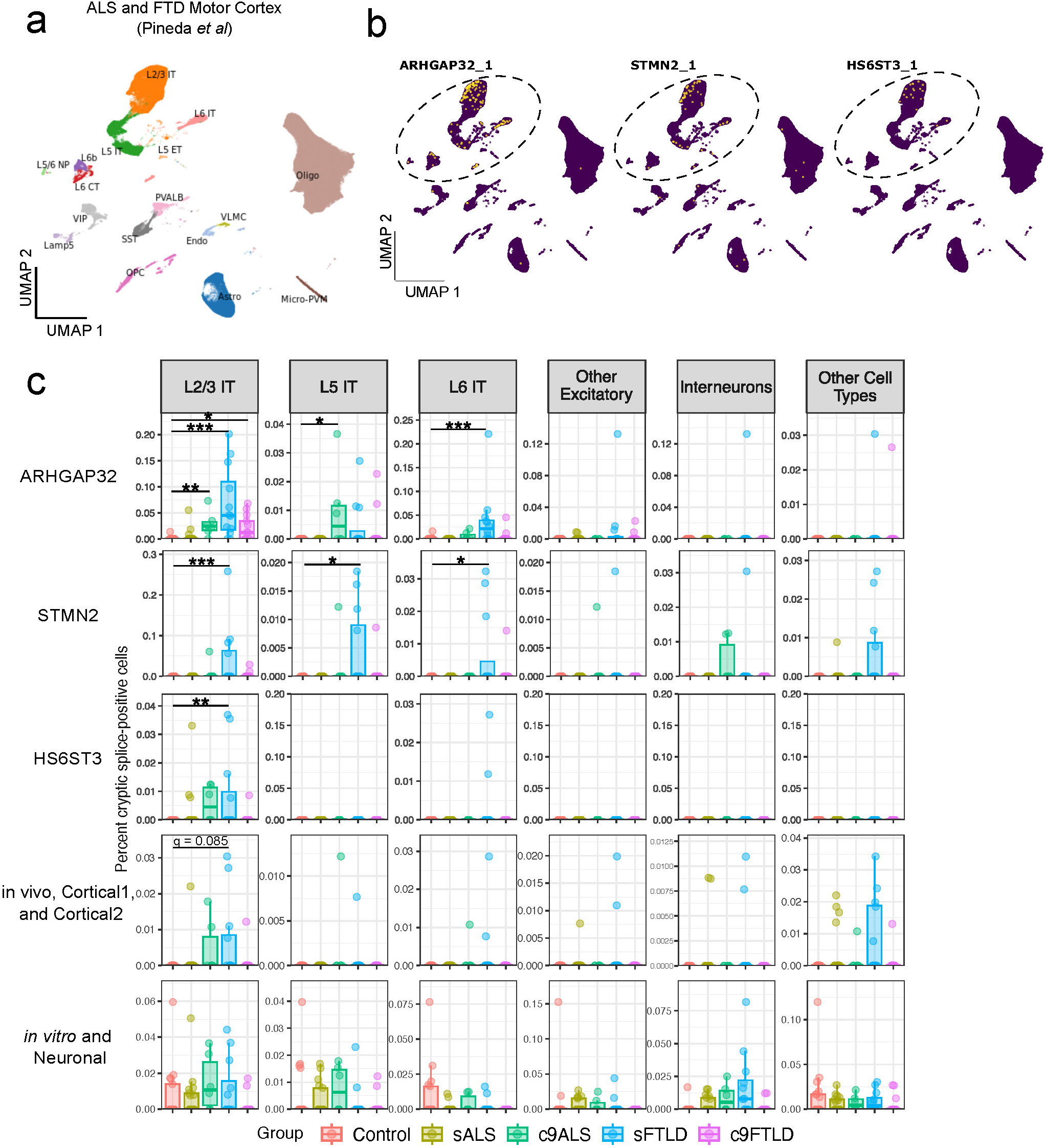
Cryptic splices are enriched in L2/3, L5, and L6 excitatory neurons in ALS and FTLD motor cortex in snRNAseq. **a** - UMAP embedding showing identified cell populations in analysis of snRNAseq data from Pineda *et al*. **b** - UMAPs showing cryptic splice-harboring cells in yellow, for splices detected at ≥10 junction reads from STMN2, cortical1, or cortical2 splice sets. Dashed circle highlights populations of excitatory neurons harboring the majority of cryptic splicing. **c** - Cellularity plots showing the percentage of cryptic-positive cells, for each cell-type, relative to the total number of detected cells (per donor). “Other excitatory” combined L5 ET, L5/6 NP, L6b, and L6 CT populations, Interneurons combined Lamp5, VIP, SST, and PVALB populations, with all other populations combined in “Other Cell Types’’. Population-wise cryptic cellularity for “Other Cell Types’’ is visualized in Online Resource 14. Each dot represents an individual donor. Significance was derived from differential abundance testing using edgeR, to account for potential composition effects. * q<0.05, **q<0.01, ***q<0.001.

In both datasets, we quantified cryptic splice junctions from CellRanger filtered BAM alignments, 41 cryptic splices were detected in Pineda (Online Resource 13: upper), with six at ≥10 reads across the whole dataset (Figure 5b). All were terminal or penultimate CEs, reflecting 3’ bias in snRNAseq data. Detection of cryptic splice junctions was greater in Pineda than in Li, likely due to 7-fold greater nuclei counts. Notably, these six cryptic splices were the most abundant in all datasets (Online Resource 13: middle and lower), indicating that cryptic splices are sparse, though consistently detected.

ARHGAP32, STMN2, and HS6ST3 cryptic splices were largely localized to L2/3, L5, and L6 excitatory neurons (Figure 5b, Online Resources 15b and 17b), and were virtually undetected in controls (Figure 5c, Online Resources 15c and 17c). Differential abundance analysis of cryptic-junction-harboring cells using edgeR, in Pineda, confirmed this specificity as statistically significant for ARHGAP32 in L2/3 (in C9-ALS, sFTLD, and C9-FTLD), L5 (C9-ALS only) and L6 excitatory neurons (in sFTLD only), for STMN2 in L2/3 neurons (in sFTLD), and for HS6ST3 (Figure 5c). Other *in vivo*, cortical1, and cortical2 splices, in aggregate, were detected in a virtually disease-specific manner (though not statistically significant), while *in vitro*-enriched splices were not disease-specific and were also not significant (Figure 5c). Non-neuronal cell-types showed sporadic and minimal inclusion of cryptic splices (Online Resources 14, 16, and 18)

Differential cryptic cell abundance analysis of Li motor and frontal cortex yielded similar results to Pineda, though only elevated STMN2 (in C9-FTLD) cryptic splicing in L6 excitatory neurons was significant, likely reflecting the greater sparsity of this dataset (Online Resources 15c and 17c). Apart from STMN2, C9-ALS samples had greater detection of cryptic splices than C9-FTLD, representing the main contrast from the Pineda analysis. ARHGAP32, HS6ST3, and other *in vivo-*enriched splices were observed virtually disease-specifically (though again not statistically significant) in L2/3 excitatory neurons in C9-ALS motor cortex.

This analysis validates ARHGAP32 and HS6ST3 as cryptic splicing targets and localizes cryptic splicing, as expected, to neurons, with apparent enrichment in intra-telencephalic projection neurons of layers 2/3, 5 and 6. The failure to detect cryptic splices in the L5 ET neurons, which would include the Betz cells expected to have TDP43 pathology in ALS, may be due to the combination of very small numbers of such cells, rarity of cryptic junctions in these datasets, and potential survivor bias in remaining Betz cells in ALS (i.e. cells with the most TDP43 pathology may already have degenerated pre-mortem). Motor cortex cryptic splicing (reaching statistical significance) appeared more prevalent in spontaneous-FTLD than ALS, if modestly. While this difference cannot be definitively confirmed here (particularly given the sparsity of snRNAseq data), it is consistent with modestly (though significantly) elevated HS6ST3, STMN2, and ARHGAP32 cryptic splice inclusion in ALS/FTD vs ALS bulk motor cortex RNAseq (Online Resource 19).

## Discussion

In this study, we meta-analyzed nearly 1,800 published RNAseq profiles of ALS and FTLD-TDP post-mortem tissue, and *in vitro* TDP43-KD models, identifying 142 cryptic splices, 68 of which had not been identified in previous CE screens. We described distinct splicing signatures across *in vivo* and *in vitro* contexts, most clearly in the form of spinal and cortical signatures which were predominantly observed in ALS and FTLD-TDP, respectively. This spinal signature did not show evidence of direct TDP43-regulation, but showed ALS-specificity and association with TDP43 pathology severity. We performed extensive computational validation of the ALS/FTD-specificity in other CNS diseases and healthy tissues by incorporating vast public bulk RNAseq databases (representing >10,000 RNAseq profiles). We validated 28 of these splices in an independent cohort of ALS lumbar spinal cord and motor cortex samples by qPCR (and three selected splices by ISH), where we confirmed context-specific cryptic splicing. Lastly, investigation of snRNAseq profiles of motor and frontal cortex demonstrated that cryptic splicing at the 3’ end of transcripts is detectable at single-nucleus resolution, and disease-enriched in L2/3, L5, and L6 excitatory neurons.

Our primary finding in this study was the discovery that cryptic splicing profiles of TDP43-proteinopathies are context-dependent, with particularly notable differences in spinal and cortical cryptic splicing signatures. In RNAseq these signatures were largely distinct, and appeared to overlap only with STMN2. RT-qPCR experiments, more sensitive than RNAseq, revealed a somewhat greater overlap between the two tissues than RNA-Seq but still confirmed many of the splicing differences(Figure 4b). The presence of many of these splices in SOD1 mutation carriers suggests that at least some of them may be TDP43-independent, and non-neuronal, reflecting a distinct physiological process. However, analysis of other datasets indicates that the spinal splices profile is still highly ALS-specific (Online Resource 7a). Furthermore, two genes harboring spinal splices have possible genetic links to the ALS/FTLD disease spectrum. CHCHD6 forms part of the MICOS complex, involved in structural maintenance of mitochondrial cristae and forms protein-complex interactions with CHCHD10, a gene which harbours autosomal dominant mutations (e,g. S59L) causative for ALS/FTD [40–42]. A recent GWAS of FTLD-TDP found a disease-associated variant linked to COL22A1 (rs146589681)[43], a splice which also shows a GT-rich region near the beginning of the cryptic exon (see <u>supplementary website</u>) potentially implicating the DNA-binding activity of TDP43[44].

Aside from tissue differences, we observed a set of six splice events exclusively *in vivo* in cortical and some spinal cord samples, but not in *in vitro* datasets. Analogous to the finding of spinal cryptic splices in SOD1 carriers, most of these splices were also included in FTLD-Tau cases, and therefore are likely due to a TDP43-independent process.

The preponderance of novel cryptic splices identified in this study reflects our investigation of *in vivo* tissue, as our analyses of *in vitro* models were concordant with those previously reported (Online Resource 1). Cryptic splicing of STMN2 and the novel cryptic splice HS6ST3, were disease-enriched in our investigation of snRNAseq datasets, particularly in cellular populations known to be enriched in TDP43 aggregates (Figure 5, Online Resources 15 and 17). Similar to another recent investigation of cryptic splicing in ALS/FTD frontal cortex, cryptic splices were sparse in snRNAseq but nonetheless detectable and enriched in disease [38]. Our localization of elevated cryptic splices in L2/3, L5, and L6 IT neurons broadly replicates this recent study by Gittings and colleagues, where the greatest STMN2 cryptic splicing was in similar layer-specific IT neuron populations [38]. TDP43 inclusions are largely localized in L2/3 of FTLD-TDP neocortex and motor cortex, extending to L6 in non-type A FTLD-TDP, consistent with our localization of FTLD cryptic splicing in L2/3, L5, and L6 IT neurons [45,46]. Cryptic splices in ALS were localized to L2/3 and L5 IT neurons (though disease-elevated with less statistical significance given the sparsity described above), which have also been reported to harbor TDP43-pathology in addition to the well-characterized pathology in Betz cells [47]. Given the capture biases in droplet-based snRNAseq[48], rarity of cryptic splice junctions, and potential for dropout in snRNAseq, the absolute detection rates in Figure 4c should be regarded as an under-estimate. Nonetheless, they do mirror observed rates of neuronal cytoplasmic inclusion burden in ALS motor cortex[49].

Notably, tissue or cell-type-specific expression of cryptic splice host genes was only a modest driver of tissue-specific cryptic splicing (Figure 3). Cryptic splices in the cortical1 splice signatures were largely in cortex-enriched genes (Figure 3a, Online Resource 9), suggesting that cortical-enriched cryptic splicing could be by way of context-enriched host gene expression. Context-specificity of other splice sets were not explained to the same degree by enriched host gene expression Another possible explanation for these differences could be tissue-or cell-specific ensembles of RNA binding proteins capable of compensating for loss of TDP43 activity at different CEs, although our analysis of CLIP-Seq binding sites (Online Resource 12) did not yield any compelling candidate regulators of the spinal splices. A more likely explanation for the context-specific spinal splices is that they reflect a regional physiological process such as gliosis Further studies should aim to address the limitations of our work. First, we did not systematically validate the TDP43-dependence of our (non-spinal splice set) cryptic splices using experimental measures. However, many were associated with metrics of TDP43 pathology and/or harboured TDP43 CLIP-peaks/putative TDP43-binding sites. Second, many cryptic splices in the spinal cord appear below the limit-of-detection by bulk-RNAseq. Future discovery efforts in spinal tissue should prioritize FACS-based enrichment of TDP43-depleted cells or nuclei to improve detection. Third, our modification of the Basescope protocol to co-detect ISH and pTDP43-IHC may have altered some cryptic ISH signal, particularly CHCHD6 which shows substantial background. Lastly, we lacked FTLD-TDP cortical tissue for validation experiments, where a cryptic splicing profile partially similar to ALS motor cortex would be expected.

In summary, our meta-analysis revealed novel cryptic splices and hitherto uncharacterized tissue, disease, and model-specific cryptic splicing signatures related to TDP43 proteinopathies. We believe that our catalogue of cryptic splices and the associated source data, companion website, and integration with public CLIPseq data and GTEx will prove a useful resource for discovery efforts. The differences in cryptic splicing between the spinal cord and cortex, ALS and FTLD-TDP, and *in vivo* and *in vitro* contexts have important implications for disease biology and biomarker development.

## Methods

### Data ingestion, QC, and raw data analysis

Publicly-available RNAseq datasets were downloaded from the Gene Expression Omnibus (GEO), Sequence Read Archive (SRA), or European Genome-Phenome Archive (EGA), as available. Dataset sources and descriptions are listed in Table 1. In all cases, raw, unprocessed FASTQ files were obtained for use with our GSNAP/HTSeqGenie-based alignment and quantification pipeline [18,19,50]. All FASTQ files were processed with FastQC [51], and files failing the *Basic Statistics* or *Per base sequence quality* measures were excluded from downstream analysis.

Bulk RNAseq data was processed using HTSeqGenie [50]. First, reads with low nucleotide quality (70% of bases with Phred quality <23) or matches to rRNA and adapter sequences were removed. The remaining reads were then aligned to the human reference genome (GRCh38.p10) using GSNAP [18,19] version ‘2013-10-10-v2’, allowing maximum of two mismatches per 75 base sequence (parameters: ‘-M 2-n 10-B 2-i 1-N 1-w 200000-E 1--pairmax-rna=200000--clip-overlap’). Transcript annotation was based on the Gencode genes database (GENCODE 27). To quantify gene expression levels, the number of reads (or concordant fragments for paired-end chemistry) mapping unambiguously to the exons of each gene was calculated.

snRNAseq data was processed using CellRanger v7.1.0 using largely default parameters. Reads were aligned to the human reference genome GRCh38, and gene expression quantification was performed using *cellranger count* (“--chemistry” was set to “threeprime” and “–include_introns” was set to true).

### Splice discovery and quantification with SGSeq

We used the SGSeq R/Bioconductor package [12] to discover and quantify alternative splicing. GRCh38 reference annotations were used to build a splice graph describing the potential splicing of each gene, in which non-overlapping exon bins are represented as nodes and splice junctions as edges (SGSeq::convertToTxFeatures). Additional splice graph features, not present in the reference annotations, were identified from the input BAM alignments (SGSeq::predictTxFeatures). Regions of the splice graph with multiple paths and shared start and/or end nodes represent *splice events*, with the particular paths representing *splice variants* (SGSeq::findSGVariants). Junction reads supporting the 5’ and 3’ ends of splice variants were quantified. The relative usage of each splice variant, *variant frequency* (VF), was calculated as a ratio of the junction reads supporting the start or end of the variant versus the sum of such reads for all variants in a splice event (SGSeq:::getSGVariantCountsFromBamFiles; e.g. figure 1B). For cases such as cassette exons with multiple junctions, the weighted mean of the variant start and end were used.

Cryptic splice events classified as alternative starts or alternative ends were inspected for internal consistency (i.e. that an upstream or downstream splice junction, respectively, was not missed), to ensure there were not indeed cassette exons. Events mis-characterized in this way were corrected by including the appropriate junction that had at least 15% the coverage of the identified cryptic junction. For mis-characterized alternative starts the new cryptic junction included must be an upstream junction terminating within the cryptic exon whereas for mis-characterized alternative ends this was a downstream junction originating within the cryptic exon.

Effects of cryptic splice inclusion on transcript sensitivity to NMD was determined by first approximating the transcripts resulting from cryptic splice inclusion. Each cryptic splice variant was inserted into each compatible canonical transcript (i.e. those containing flanking exons with splice sites matching the cryptic splice junctions) for the respective gene, and resulting cryptic transcript sequences were determined. Cryptic transcripts which introduced premature stop codons more than 50bp upstream of the previous splice junction were classified as NMD candidates[52]. Splice variants with non-uniform splice effects (e.g. some cryptic transcripts classified as NMD candidates and some not), were classified as “mixed”.

SGSeq analyses were performed separately within datasets to improve cryptic splice detection across varying contexts, with four combined analyses performed across similar contexts to increase sample sizes (indicated inOnline Resource 21). This reflected combining all spinal cord, NYGC frontal/temporal cortex, and *in vitro* contexts into respective SGSeq analyses. Parameters controlling the inclusion of splice graph features were modified slightly to optimize detection across datasets with differing sequencing depths and levels of intronic reads. Namely *β* (the FPKM threshold for splice junction inclusion) and the *min_samp* parameter (minimum number of samples in which a splice graph feature must be observed to be included) were modified as described inOnline Resource 21. Splice variant nomenclature has the format: “GeneID_EventNumber_VariantNumber/TotalVariants_EventType” (e.g. STMN2_2_1/2_AE).

### Differential splicing analyses with DEXseq

Variant count quantifications were used as input to DEXseq to test for differential usage of splice variants within an event, similar to a previously described approach [53]. Splice variants were included if supported by at least 10 reads, in at least *n* samples (n=2 for in vitro models, n=4 for Liu and RiMOD, and n=8 for all other analyses). Any events with a single variant remaining (i.e. singletons) were excluded from analysis, as they were considered effectively constitutively spliced. After filtering, DEXseq analysis proceeded as previously described [53]: normalizing for sequencing depth (DEXSeq::estimateSizeFactors), estimating transcriptome-wide dispersion (DEXSeq::estimateDispersions), and testing for differential variant usage within an event (DEXSeq::testForDEU).

Differential contrasts were performed between ALS and controls in the hippocampus, cerebellum, spinal cord, motor cortex, frontal cortex, and temporal cortex, between FTLD-TDP and controls in the frontal cortex, temporal cortex, and RiMOD datasets, between TDP43-negative nuclei and TDP43-positive nuclei from the Liu dataset, and between TDP43-KD siRNA treatments and scrambled siRNA controls in *in vitro* contexts. Statistical significance was set at a Benjamini-Hochberg[54] FDR of 5% and a ΔVF of 0.025. Cryptic splices were identified as significantly differentially spliced variants with a control group mean or median VF of 0.05 or less.

Variant frequencies within an event sum to 1, thus inclusion of all variants in DEXseq analysis would result in excessive multiple testing correction. In our DEXseq analysis, we excluded the first variant from each event to address this, despite this resulting in some disease-elevated cryptic variants being excluded (though significance is identical for the canonical variant with 1-VF). For all significantly differentially spliced variants, the disease-elevated variants were confirmed *post-hoc*. Cryptic splice variants were consolidated across all analyses by de-duplicating independently identified variants with identical cryptic junctions.

Variant frequency Z-scaling (used only in visualization of the figure 2A heatmap) was used to more clearly compare variant frequency across datasets with vast differences in effect size. Variant frequency was normalized to the mean and standard deviation of the respective control group, with a pseudocount included for the standard deviation to avoid dividing by zero when a splice was completely undetected in controls in a particular context. 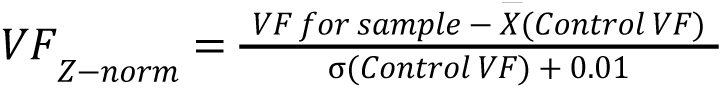

### Post-mortem lumbar spinal cord tissue

All human tissue samples were supplied by Discovery Life Sciences and Fidelis Research. Each supplier received IRB approval of research, appropriate informed consent of all subjects contributing biological materials, and all other authorizations, consents, or permissions as necessary for the transfer and use of the biological materials for research at Genentech.

### pTDP43 IHC, imaging, and quantification

Frozen spinal cord and motor cortex samples embedded in OCT were sectioned transversely at 8 um thickness, and IHC was performed using the Ventana Discovery Ultra platform (Ventana Medical Systems). Sections were fixed for 30 minutes in neutral buffered formalin (VWR; Radnor, PA: cat# 16004-122), heat antigen retrieved with Ventana CC1 for 8 minutes (cat. no. 950-500, Ventana Medical Systems), blocked for 4 min (DISCOVERY Inhibitor; Ventana, 760-4840), and incubated with 1:2000 mouse anti-human TDP43 pS409/410 (Cosmo Bio TIP-PTD-M01A; clone 11-9) for 60 minutes. Signal was detected with Ventana Mouse Omnimap anti-mouse HRP (Ventana Medical Systems; Tucson, AZ: cat# 760-43100) for 16 minutes, and labeled with Ventana Discovery Purple (Ventana: cat# 760-229) incubation for 12 minutes. The sections were counterstained with Hematoxylin II (Ventana: cat# 790-2208), dehydrated, and cover-slipped. Negative control IHC was performed using naive mouse IgG1 antibody.

Semiquantitative analysis of pTDP43 IHC was performed by a pathologist (JH) blinded to the cryptic splicing quantifications and donor diagnoses, and was scored as follows: 0, No pTDP43; 1, Minimal pTDP43 aggregates in rare cells; 2, Moderate pTDP43 aggregates in scattered cells; 3, Severe pTDP43 aggregates in frequent cells. No healthy control samples had pTDP43 aggregates in the sections examined.

### Cryptic Basescope ISH + pTDP43

BaseScope ISH was combined with IHC to enable the co-detection of cryptic spliced mRNA and pTDP-43 protein on the same slide using a modified protease-free workflow designed to preserve the protease-sensitive pTDP-43 antigen. Frozen spinal cord samples embedded in OCT were sectioned transversely at 8 µm thickness. The sections were fixed in Neutral Buffered Formalin for 90 minutes (cat. no. 16004-122, VWR). Antigen retrieval was performed using BOND Epitope Retrieval Solution 2 (cat. no. AR9640, Leica Biosystems) for 10 minutes and RNAscope 2.5 LS PretreatPro (RNAscope 2.5 LS Pro Reagents Kit, cat. no. 322020, Advanced Cell Diagnostics) for 30 minutes.

The ISH procedure was carried out using the BaseScope LS Reagent Kit (cat. no. 323600, Advanced Cell Diagnostics), the BOND Polymer RED Detection Kit (cat. no. DS9390, Leica Biosystems), and custom 1ZZ BaseScope probes (BA-Hs-STMN2-E1cE2-C1, cat. no. 1561798-C1; BA-Hs-CHCHD6-cE6E7-C1, cat. no.

1561808-C1; BA-Hs-KALRN-cE56E57-C1, cat. no. 1559518-C1, Advanced Cell Diagnostics). The standard BaseScope ISH protocol was followed with one modification: BaseScope LS AMP7 was replaced with RNAscope 2.5 LS AMP Pro (RNAscope 2.5 LS Pro Reagents Kit, cat. no. 322020, Advanced Cell Diagnostics). The sections were coverslipped, and ISH staining was imaged using the NanoZoomer S60 Digital Slide Scanner.

After imaging, the coverslips were removed, and the second part of the RNA-protein co-detection—pTDP-43 IHC—was performed on the BOND Rx automated staining platform (Leica Biosystems). Heat-induced epitope retrieval was repeated with BOND Epitope Retrieval Solution 2 (cat. no. AR9640, Leica Biosystems) for 10 minutes, followed by blocking with 3% hydrogen peroxide solution for 4 minutes (cat. no. H1009, Millipore Sigma). Sections were incubated with 1:2000 mouse anti-human TDP-43 pS409/410 (Cosmo Bio TIP-PTD-M01A; clone 11-9) for 60 minutes. The IHC signal was detected using PowerVision Poly-HRP Anti-Mouse IgG (cat. no. PV6114, Leica Biosystems) for 15 minutes and visualized with a green chromogen (cat. no. DC9913, Leica Biosystems). Sections were counterstained with Hematoxylin (cat. no. AR9915, Leica Biosystems), coverslipped, and imaged using the NanoZoomer S60 Digital Slide Scanner.

STMN2 cryptic ISH signal was clearly cellular (overlaid on pTDP43 signal and), and the number of STMN2 cryptic ISH-positive motor neurons (identified by morphology and location) in the anterior horns of each sample was counted and compared between control and ALS groups using unpaired t-tests. CHCHD6 cryptic ISH signal was diffuse, not consistently associated with cells/nuclei, and of non-uniform size suggesting substantial background signal.

### RT-qPCR validation of cryptic splices

Spinal cord or motor cortex tissue was homogenized using Bertin-Precellys homogenizer in Buffer RLT (Qiagen) supplemented with beta-mercaptoethanol per manufacturer recommendations. RNA was prepared using the RNeasy Plus RNA isolation kit (Qiagen) and quantified using Tapestation High sensitivity RNA kit (Agilent).

Levels of RNA were normalized for cDNA synthesis with SuperScript IV VILO master mix (Thermo Fisher). Transcripts were quantified by real-time quantitative PCR on a Quantstudio 7 Real-Time PCR instrument (Thermo Fisher). Probes for 67 cryptic splice variants were designed using Primer3 (Online Resource 22) [55], and synthesized by Integrated DNA Technologies. For each sample, mRNA abundance was normalized to GAPDH (Hs02786624_g1, Thermo) and HPRT1 (Hs02800695_m1,Thermo) transcripts. Triplicate PCRs were performed on each sample of cDNA and the mean value of each sample was used in statistical analysis.

We quantified relative splice variant expression using the ΔΔCt method: 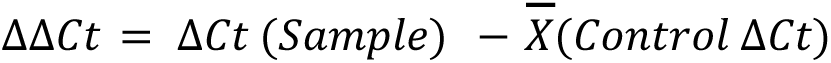

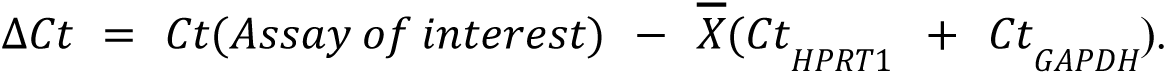 For ΔΔCt values in Figure 4A, cycle thresholds (Cts) for unamplified assays were imputed as (1 + 𝐿𝑎𝑟𝑔𝑒𝑠𝑡 𝐶𝑡 𝑓𝑜𝑟 𝑎𝑠𝑠𝑎𝑦), no imputation was performed for ΔCt values (i.e. not normalized to experimental controls) shown in any other figure. ΔΔCt values were compared between ALS and control samples by unpaired t-test, significance was set at p < 0.05.

### Bulk-RNAseq of lumbar spinal cord samples from ALS and controls

Total RNA was quantified with Qubit RNA HS Assay Kit (Thermo Fisher Scientific) and quality was assessed using RNA ScreenTape on TapeStation 4200 (Agilent Technologies). For sequencing library generation, the Truseq Stranded mRNA kit (Illumina) was used with an input of 100-1000 nanograms of total RNA. Libraries were quantified with Qubit dsDNA HS Assay Kit (Thermo Fisher Scientific) and the average library size was determined using D1000 ScreenTape on TapeStation 4200 (Agilent Technologies). Libraries were pooled and sequenced on the NovaSeq 6000 platform (Illumina) to generate 50 million paired-end 100bp reads for each sample. Alignment and splice variant quantification were performed as described above.

### snRNAseq QC, analysis, and cell-type annotation

snRNA-seq data from Pineda *et al* (Motor Cortex), Li *et al* (Frontal Cortex), and Li *et al* (Motor Cortex) were analyzed separately, using the scuttle, scran, and scater packages [36,37,56,57]. snRNA-seq quality control metrics (total number of unique molecular identifiers (UMIs), number of detected genes, and mitochondrial read percentage) were computed for each nucleus, and sample-wise outlier-based filtering was performed to remove nuclei with any metric >3 median absolute deviations from the sample median. After initial analysis, both cortical [58]regions of the Li dataset contained a cluster of low-UMI, high mitochondrial UMI percentage, poorly annotated nuclei which were filtered by removing nuclei with fewer than 1800 UMIs or greater than 10% mitochondrial UMIs (followed by re-analysis). UMI counts were size-factor normalized within each sequencing batch, scaled across batches to match the batch with the lowest coverage, then log-transformed. Principal component analysis was performed on the 2000 most variable genes within each dataset. Batch correction was performed through mutual nearest-neighbor (MNN) integration across donors (Pineda) or biological sex (Li) on 30 components [39]. 25 MNN-corrected components were used as input to shared nearest-neighbor clustering, and further dimension reduction by uniform manifold approximation and projection (UMAP).

Cell-type annotation was performed using SingleR[59] and the Allen Brain human M1 cell-type atlas (https://portal.brain-map.org/atlases-and-data/RNAseq/human-m1-10x) as a reference. SingleR annotations were confirmed by cross-referencing with population enrichment of canonical cell-type and laminar markers: SLC17A7 (excitatory neurons), CUX2, RORB, THEMIS, FEZF2 (superficial to deep layers), GAD1, GAD2, SST, PVALB, VIP (interneurons), MBP, MAG (oligodendrocytes), PDGFRA (oligodendrocyte precursor cells), TMEM117, IBA1 (microglia), GFAP, AQP4 (astrocytes). Doublet detection was performed using scDblFInder [60], and clusters enriched in high doublet scores which exhibited expression of cell-type marker genes from different cell-types were removed from further analysis.

### Splice junction quantification and differential abundance

To identify cryptic splice junctions within single-nuclei, we used the Bioconductor GenomicAlignments and Rsamtools package to extract reads from CellRanger BAM alignments with split alignments covering cryptic splice junctions using the Rsamtools::ScanBamParam and GenomicAlignments::readGAlignments functions, for each splice. Reads were then filtered to confirm that only those compatible with the cryptic splice junction, and containing coverage in flanking cryptic or canonical exons, were included. Cryptic junction read counts were then mapped to the corresponding nucleus via sample identifiers and cell barcode tags. Given the low dynamic range in junction counts per nucleus (Online Resource 13), downstream analyses focused on the sample-wise proportion of cryptic splice-positive nuclei. For differential abundance analyses, cryptic splice-harboring cells were categorized as an additional cell-type (e.g. “cryptic STMN2 L2/3 neurons”) and included with non-cryptic cell-types.

Sample-wise abundances of each cell-type were then compared across diagnostic groups and corrected for multiple comparisons using edgeR [61,62]. Significance was set at q < 0.05.

### Data Availability

Previously published bulk and snRNAseq data sources are available in Table 1. We have deposited the raw bulk RNAseq data from our new cohort of 10 control and 10 ALS lumbar spinal cord samples to EGA (EGAS50000000575). Cryptic splice variant coordinates and quantifications are available inOnline Resource 20, pTDP43 IHC data is available in Table 2, and RT-qPCR data is available in Online Resource 23.

## Supporting information

Online Resource 23

Online Resource 22

Online Resource 21

Online Resource 20

Online Resource

