## Supplementary material for "Divergent TDP43-regulated and TDP43-independent cryptic splicing in the cortex and spinal cord": Online Resource

**Table of Contents**

Supplementary Figures (This document)

**Online Resource 1.** Comparison of SGSeq-DEXseq splicing analysis in TDP43-KD NGN2-iPSCs to published MAJIQ analysis**.**

**Online Resource 2.** Dotplot of cryptic splice inclusion

**Online Resource 3.** Examples of context-specific splices for the six identified splice sets.00

**Online Resource 4** A subset of cryptic splices identified from in vitro models are either ubiquitous or not associated with disease.

**Online Resource 5.** Inclusion of STMN2 cryptic splice, with spinal and *in vivo* splice sets in spinal cord and motor cortex, with SOD1-ALS.

**Online Resource 6**. Spinal-cord splicing profile is disease-enriched in independent spinal cord RNAseq datasets.

**Online Resource 7.** Inclusion of literature-reported cryptic splices, and spinal splices, across non ALS/FTD degeneration contexts and GTEx.

**Online Resource 8.** Expression of cryptic splice-harbouring genes across Spinal Cord and Frontal, Motor, and Temporal cortex in the Human Brain Cell Atlas.

**Online Resource 9.** GT-repeats and TDP43 CLIP-seq peaks relative to cryptic exons of all cryptic splices.

**Online Resource 10.** Exploring trans-regulatory RBP factors that may explain tissue-specific enrichment of the spinal splices.

**Online Resource 11.** Relationships between cryptic splice inclusion and gene expression.

**Online Resource 12.** Detection of STMN2 and CHCHD6 cryptic splices in a novel spinal cord RNAseq cohort.

**Online Resource 13.** Distribution of cryptic splice junction counts across all cells in snRNAseq datasets.

**Online Resource 14.** Cryptic cellularity plots of all cell-types identified in analysis of Pineda *et al* dataset.

**Online Resource 15.** Cryptic splices are enriched in L6 excitatory neurons in FTLD (Li *et al* motor cortex) snRNAseq.

**Online Resource 16.** Cryptic cellularity plots of all cell-types identified in analysis of Li *et al* motor cortex.

**Online Resource 17.** Cryptic splices are enriched in L6 excitatory neurons in FTLD (Li et al frontal cortex) snRNAseq.

**Online Resource 18.** Cryptic cellularity plots of all cell-types identified in analysis of Li *et al* frontal cortex.

**Online Resource 19.** Variant frequency plots of ARHGARP32, STMN2, and HS6ST3 cryptic splices in motor cortex bulk RNAseq.

Supplementary Tables

**Online Resource 20.** Cryptic splice metadata, splice variant structures, and quantifications across all samples included in the meta-analysis.

**Online Resource 21.** Analytical details and parameters used in SGSeq analyses of each bulk RNAseq dataset.

**Online Resource 22.** RT-qPCR primer sequences used to quantify cryptic and canonical splice variants.

**Online Resource 23.** Raw qPCR data.

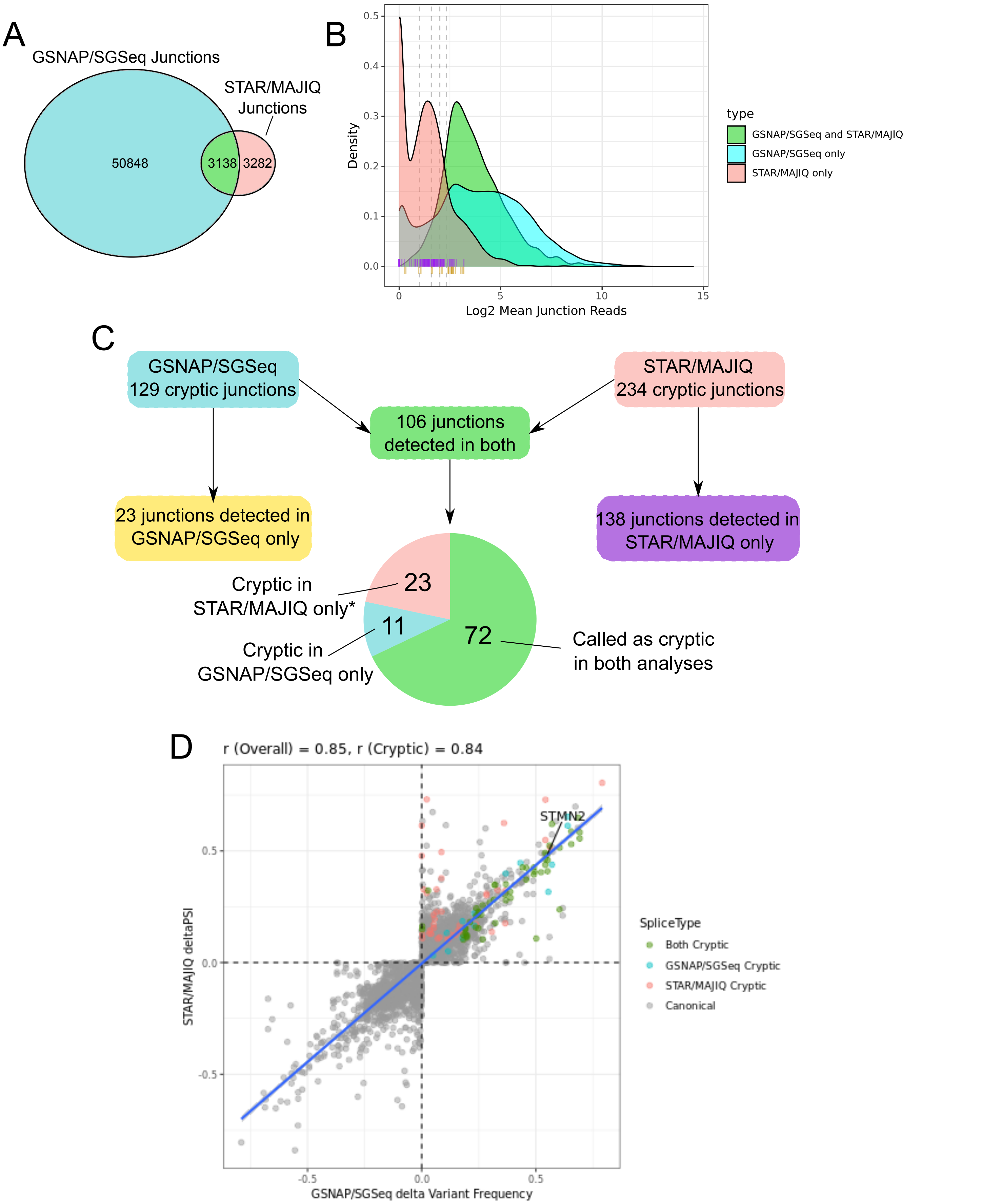


Online Resource 1. Comparison of SGSeq-DEXseq splicing analysis in TDP43-KD NGN2-iPSCs to published MAJIQ analysis. A - Comparison of all splice junctions detected in the STAR/MAJIQ analysis of TDP43-KD in NGN2-expressing iPS-neurons from Brown et al., Nature (2022) (data source: Supplementary Table 1), and the GSNAP/SGSeq-based analysis performed in the present study. The ~8x greater number of junctions observed in SGSeq largely reflects additional canonical splicing not captured by STAR/MAJIQ. B - Mean junction reads across all samples for STAR/MAJIQ-specific junctions (salmon), GSNAP/SGSeq junctions (cyan), and shared GSNAP/SGSeq-STAR/MAJIQ junctions (green). Purple lines (n=138) indicate the mean junction read depth supporting cryptic splices which were STAR/MAJIQ-specific, and yellow lines (n=23) indicate the same for GSNAP/SGSeq-specific cryptic junctions. Dashed lines are reference lines indicating 2, 3, 4, and 5 mean reads. C - Mapping of cryptic splice junctions identified in STAR/MAJIQ to those in GSNAP/SGSeq, yellow and purple boxes represent the same lines visualized in Online Resource 1B. The green box indicates cryptic splice junctions commonly identified in both analyses, with the corresponding pie chart describing the proportion of these which were called as cryptic in the two analyses. The splices in the pie chart correspond to the coloured points in Online Resource 1D. *Note - all 23 of these junctions were indeed identified in GSNAP/SGSeq analyses, but were filtered from DEXseq analyses due to low numbers of junction reads. D - Comparison of computed effect sizes for all splice junctions shared between GNSAP/SGSeq and STAR/MAJIQ analyses. Effect sizes were highly correlated, both overall (i.e. largely reflecting canonical splicing), and when subsetting cryptic splices only, indicating a broad concordance of the two analytical approaches.

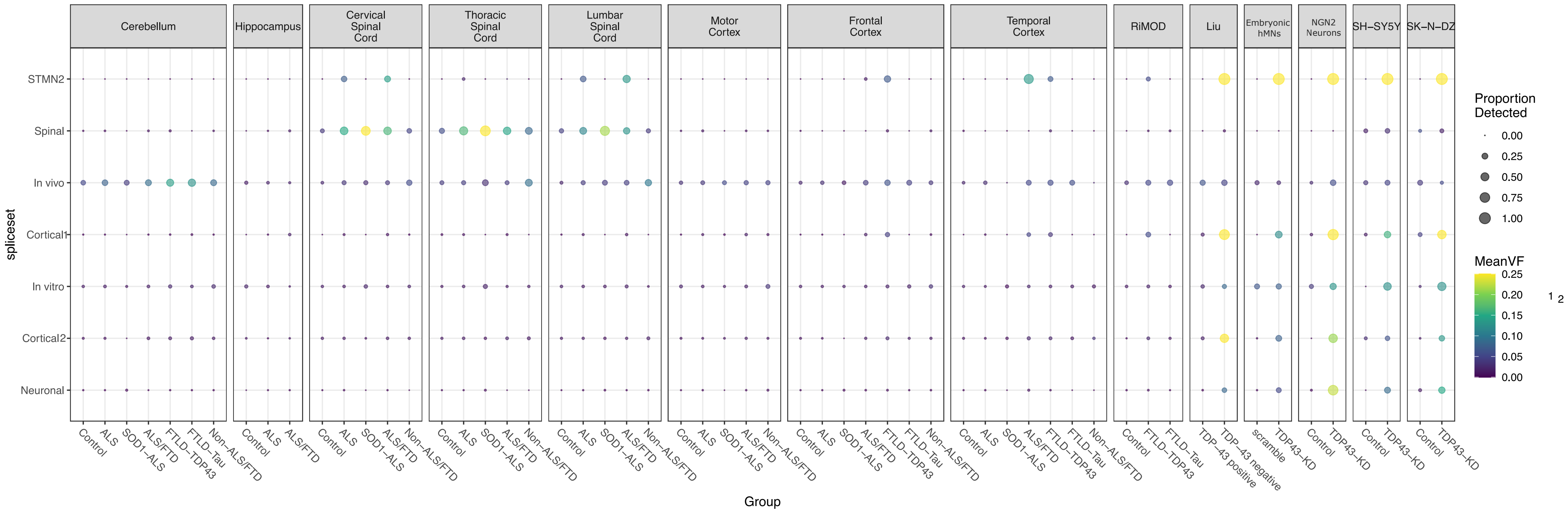


**Online Resource 2. Dotplot of cryptic splice inclusion.** Dot-plot style visualization of the cryptic splices shown in figure 2. Cryptic splices were aggregated by splice set, and their percent detection (non-0) in each group, in each dataset is shown by the size of the dot. Dot colour reflects mean variant frequency across splices, with a ceiling applied at VF=0.25 to account for extreme differences seen in TDP43-KD models. Note, the strong percent detection and variant frequency of spinal splices in SOD1-ALS subjects is exacerbated by these samples representing a maximum of 3 donors per region profiled (see figure 1A for comparable sample sizes in other groups within the NYGC cohort).

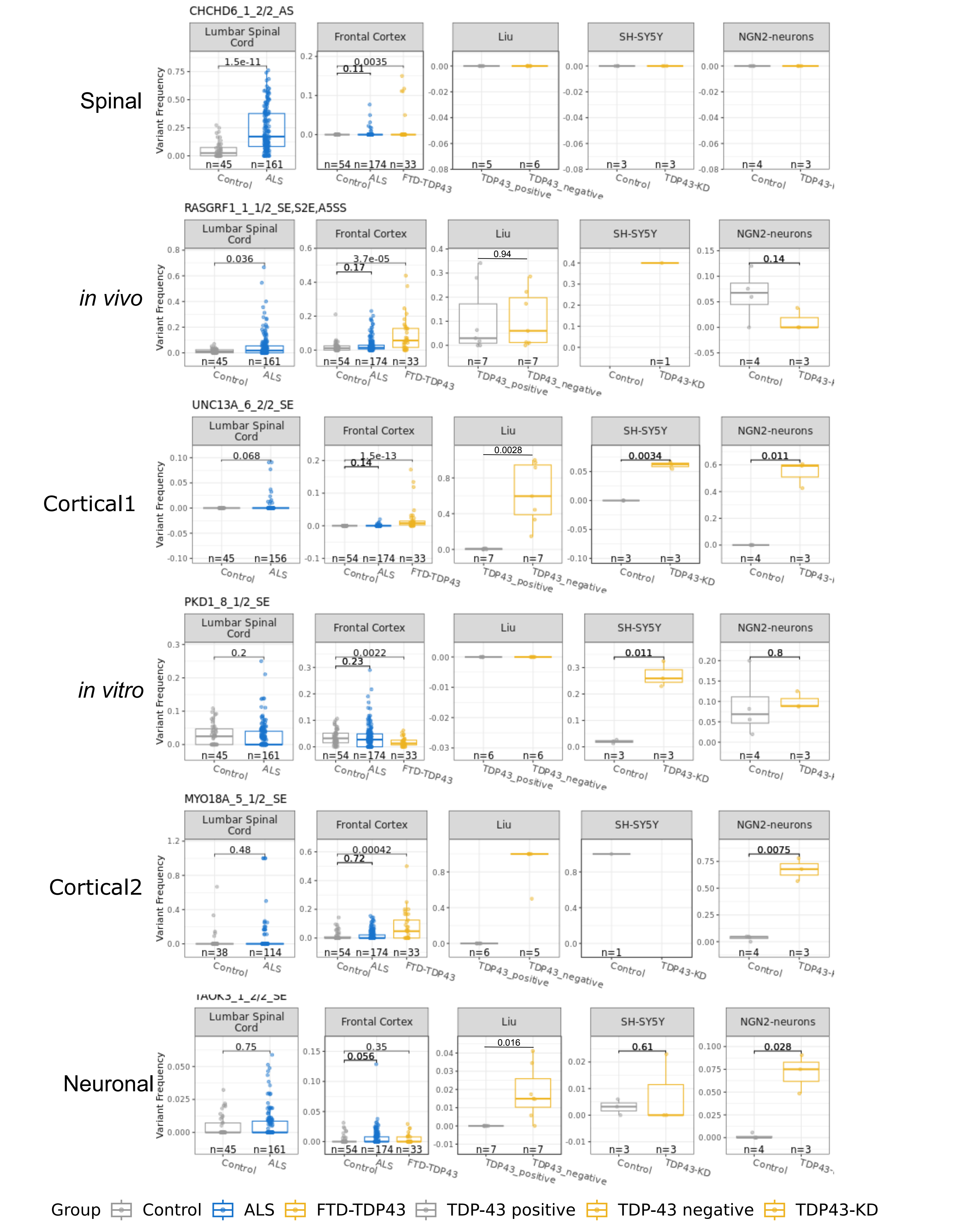


Online Resource 3. Examples of context-specific splices for the six identified splice sets. Boxplots showing variant frequency of 6 cryptic splices, representing each of the splicing signatures identified in Figure 2A. Lumbar spinal cord and frontal cortex were compared using Mann Whitney U tests, Liu using paired-samples t-tests, and SH-SY5Y and NGN2-neurons compared using unpaired t-tests. Parametric statistics were used for the latter three contexts given the low sample size and abundance of ties which undermine rank-based nonparametric tests.

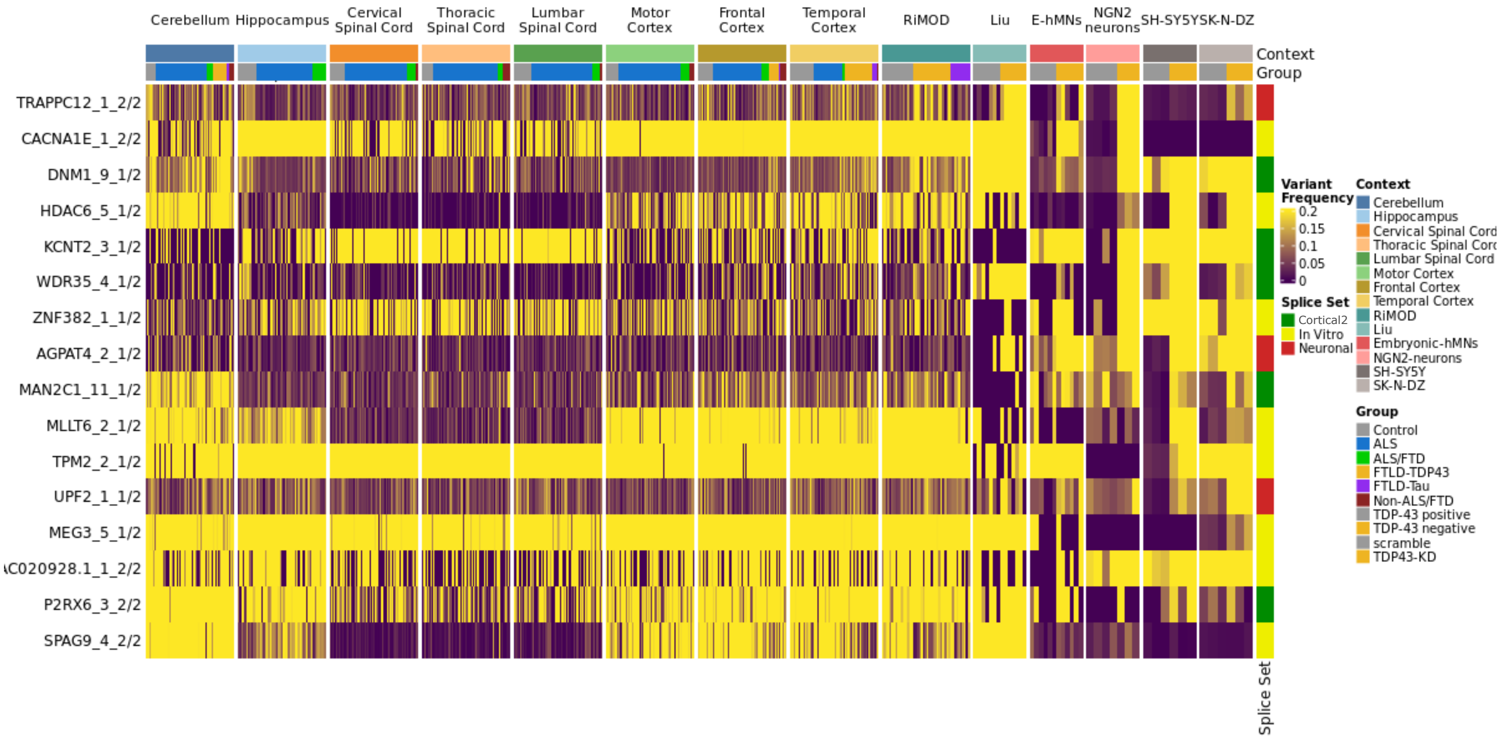


Online Resource 4. A subset of cryptic splices identified from in vitro models are either ubiquitous or not associated with disease. Heatmap showing variant frequency of 16 cryptic splices which have mean variant frequencies > 0.1 across in vivo contexts. 5 were members of the Cortical2 splice set, and 11 members of the in vitro or neuronal (largely driven by in vitro datasets, aside from the Liu et al study), though all were discovered within analyses of in vitro models.

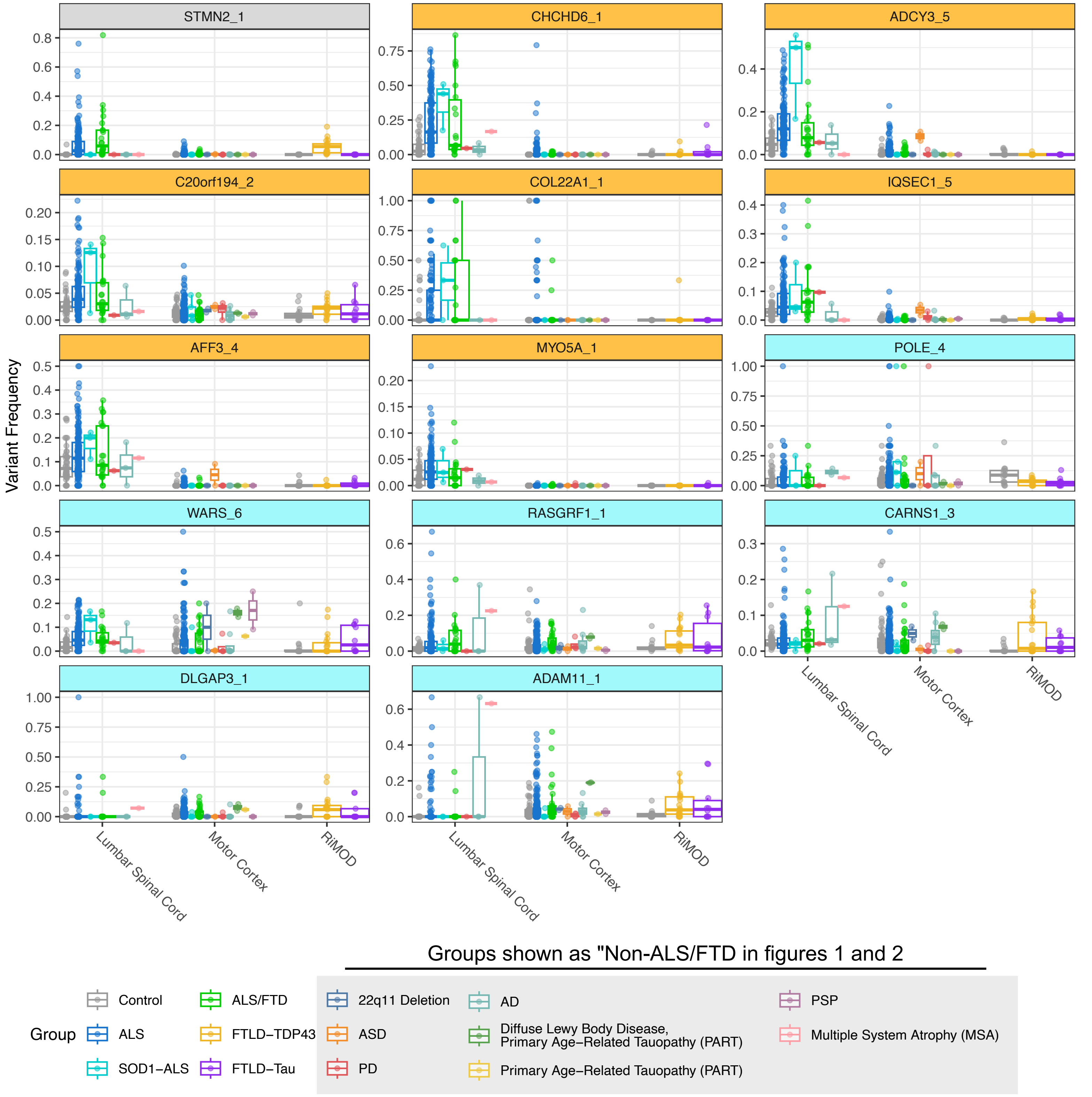


Online Resource 5. Inclusion of STMN2 cryptic splice, with spinal and in vivo splice sets in spinal cord and motor cortex, with SOD1-ALS. Boxplots showing variant frequency of splices in the spinal (orange) and in vivo (cyan) splice sets with STMN2 as a positive control comparison for TDP43-regulation. Diagnostic groups shown as “Non-ALS/FTD” in figures 1 and 2 are captured in the groups from q2211 deletion to multiple system atrophy (indicated by a grey box in the legend).. SOD1-ALS subjects show inclusion of spinal cryptic splices comparable to non-SOD1 ALS, whereas the spinal and in vivo splices largely do not (with the exception of WARS_6 in the spinal cord and POLE_4 in the motor cortex). Likewise, these in vivo splices are included in FTLD-Tau subjects,

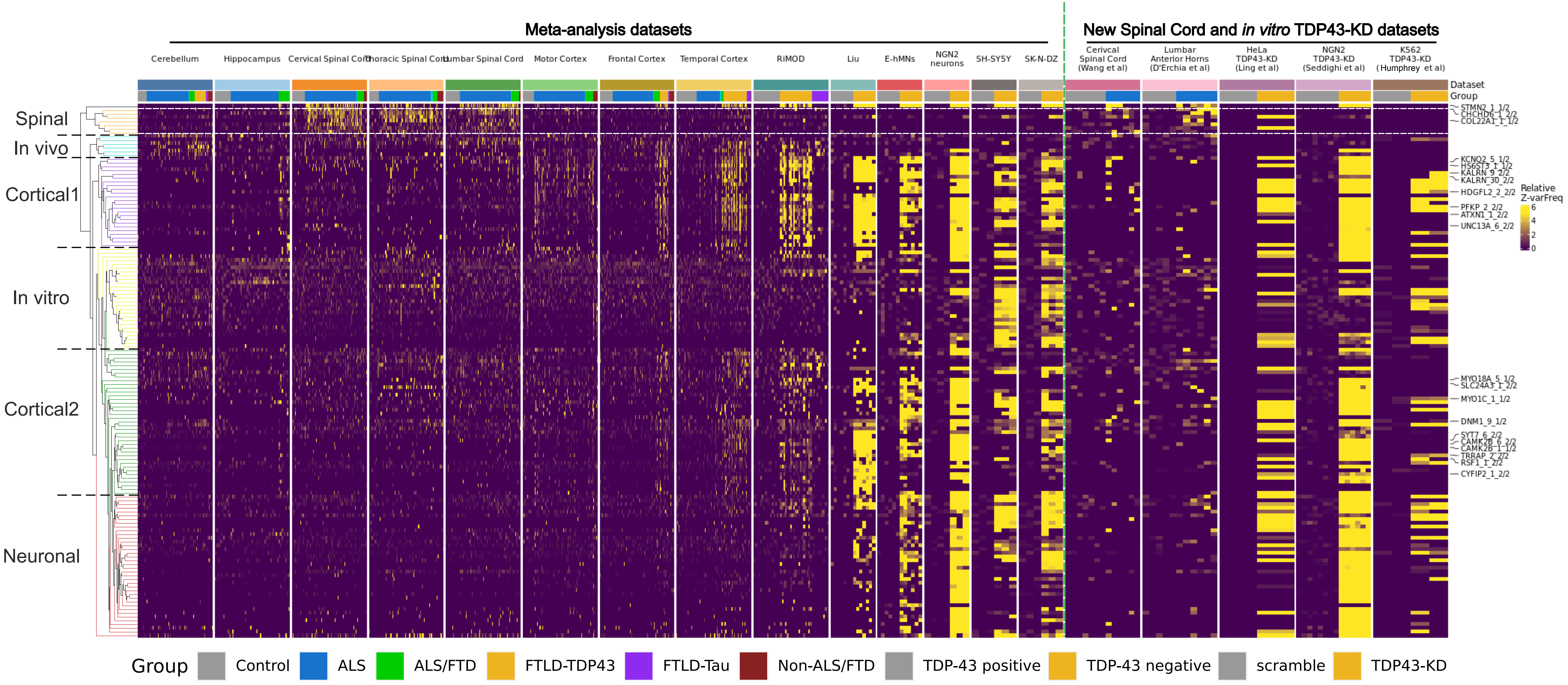


Online Resource 6. Spinal-cord splicing profile is disease-enriched in independent spinal cord RNAseq datasets. Heatmap showing all cryptic splices, as quantified in Figure 2. Meta-analysis studies shown to the left of the green dashed vertical line, and additional studies shown to the right. The spinal splice set is called out by horizontal white lines across the heatmap, demonstrating the ALS-enrichment of these splices (and STMN2) in the cervical spinal cord, and lumbar anterior horns, of independent patient cohorts, distinct from the NYGC. Additionally, three additional, independent, TDP43-KD experiments in HeLa, NGN2-iNeurons, and K562 cells show broad re-capitulation of the in vitro cryptic splicing patterns shown in the original meta-analysis. Notably, STMN2 cryptic splicing is absent from HeLa cells, consistent with the original manuscript of Ling et al, Science (2015) in which it was not reported.

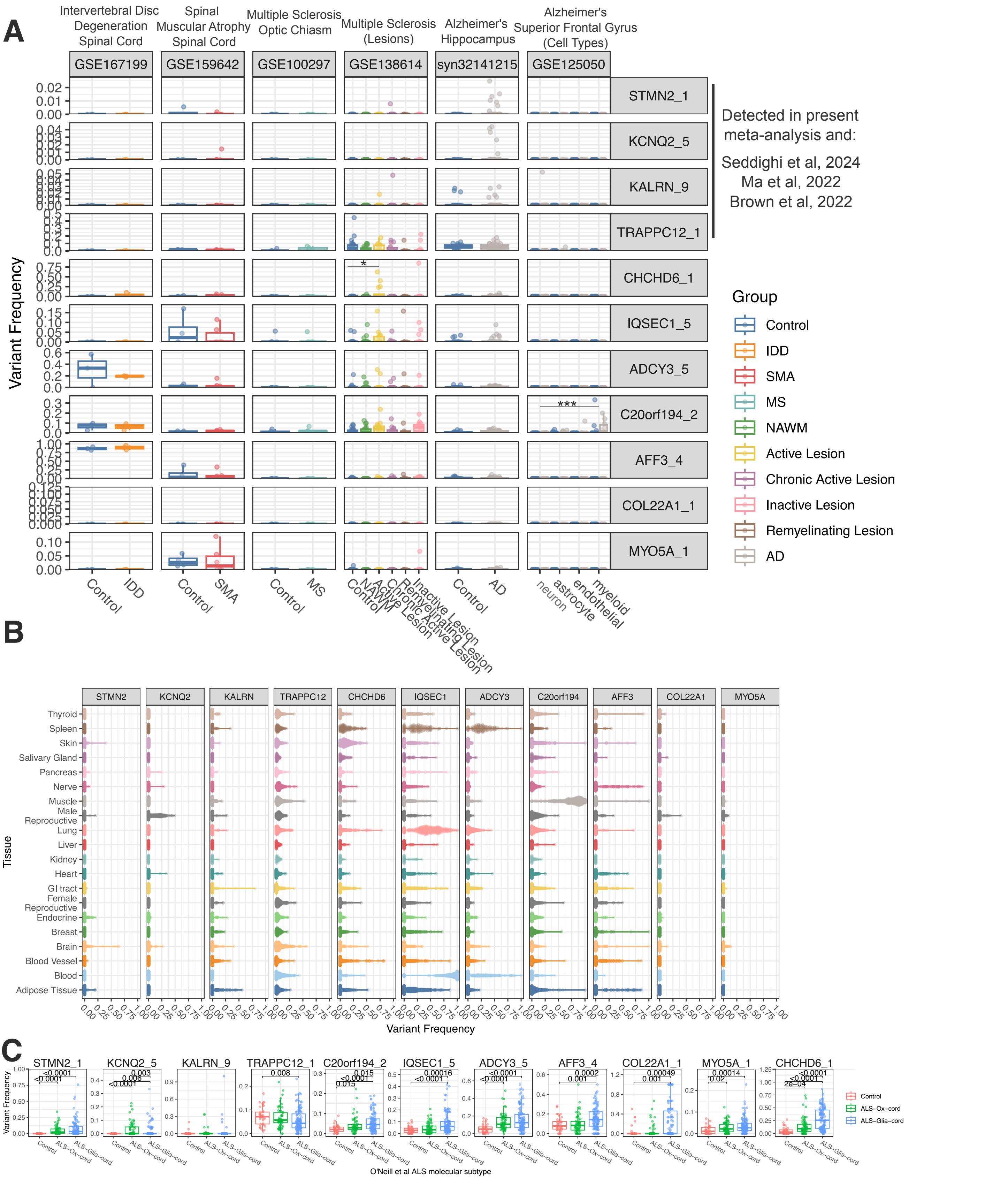


Online Resource 7. Inclusion of literature-reported cryptic splices, and spinal splices, across non ALS/FTD degeneration contexts and GTEx. A - Boxplots of previously-reported cryptic splices (STMN2, KCNQ2, KALRN, and TRAPPC12), and the spinal cryptic splices, across six public studies which performed bulk RNAseq in spinal cord injury/neurodegenerative contexts. Cryptic variant frequency was calculated for the above events, and compared across groups with unpaired t-tests (except for between cell-type comparisons of GSE125050 which had paired samples from the same donors). Broadly, previously-reported cryptic splices were not significantly elevated in any disease/injury setting compared to its respective control. TRAPPC12 did show appreciable inclusion (many samples with cryptic variant frequency > 0.05 in MS lesions, AD hippocampus, and controls). Spinal splices showed a similar minimal level of detection, with the exception of ADCY3 amad AFF3 in IDD/control spinal cord, and sporadic inclusion of CHHD6, IQSEC1, ADCY3, and C20orf194 in MS lesions. CHCHD6 cryptic splicing was nominally significantly elevated in MS active lesions versus control white matter, and C20orf194 cryptic splicing was elevated in sorted myeloid cells in AD/Controls (irrespective of disease). Notably, COL22A1 cryptic splicing was not observed in any of these contexts. These findings suggest that the spinal splices are broadly ALS-specific (or enriched), and are not a part of a more general pan-disease neurodegenerative process. * $p < 0.05$, *** $p < 0.001$. B - Violin plots of cryptic splice inclusion in the Genotype-Tissue Expression (GTEx) project, consisting of ~6,000 bulk RNAseq profiles across the human body, from ~1,000 healthy donors. Samples were removed, within each cryptic splice, if they contained less than 5 splice junction reads across the cryptic and canonical junctions combined (i.e. the splice event as a whole was not confidently detected). STMN2, KCNQ2, and KALRN show virtually zero inclusion across the body, with the exception of some KCNQ2 cryptic splicing in male reproductive tissue, and KALRN in adipose tissue. TRAPPC12 shows low-level, but detectable, inclusion across most regions of the body - suggesting it is either weakly TDP43-dependent and/or regulated by other factors. The spinal splices showed variable inclusion across healthy tissues, with CHCHD6 detectable in spleen and skin, IQSEC1 abundant in spleen, lung, and blood, ADYC3 in spleen and blood, C20orf194 in muscle and low-level inclusion in other tissues, and AFF3 was sporadically included in most tissues to varying extents. Notably, COL22A1 and MYO5A were virtually absent from all samples, similar to STMN2. C - Cryptic splice variant frequency in lumbar spinal cord samples, with ALS molecular subtypes as defined in O'Neill et al, 2025 (Cell Reports), which largely fall into glial/inflammatory (ALS-Glia-cord) and oxidative stress-related (ALS-Ox-cord) phenotypes. Note STMN2 was elevated in both subtypes and KCNQ2 was elevated in ALS-Ox more than ALS-Glia. Additionally, all spinal cryptic splices were elevated in ALS-Glia, and often to a significantly greater extent than in ALS-Ox.

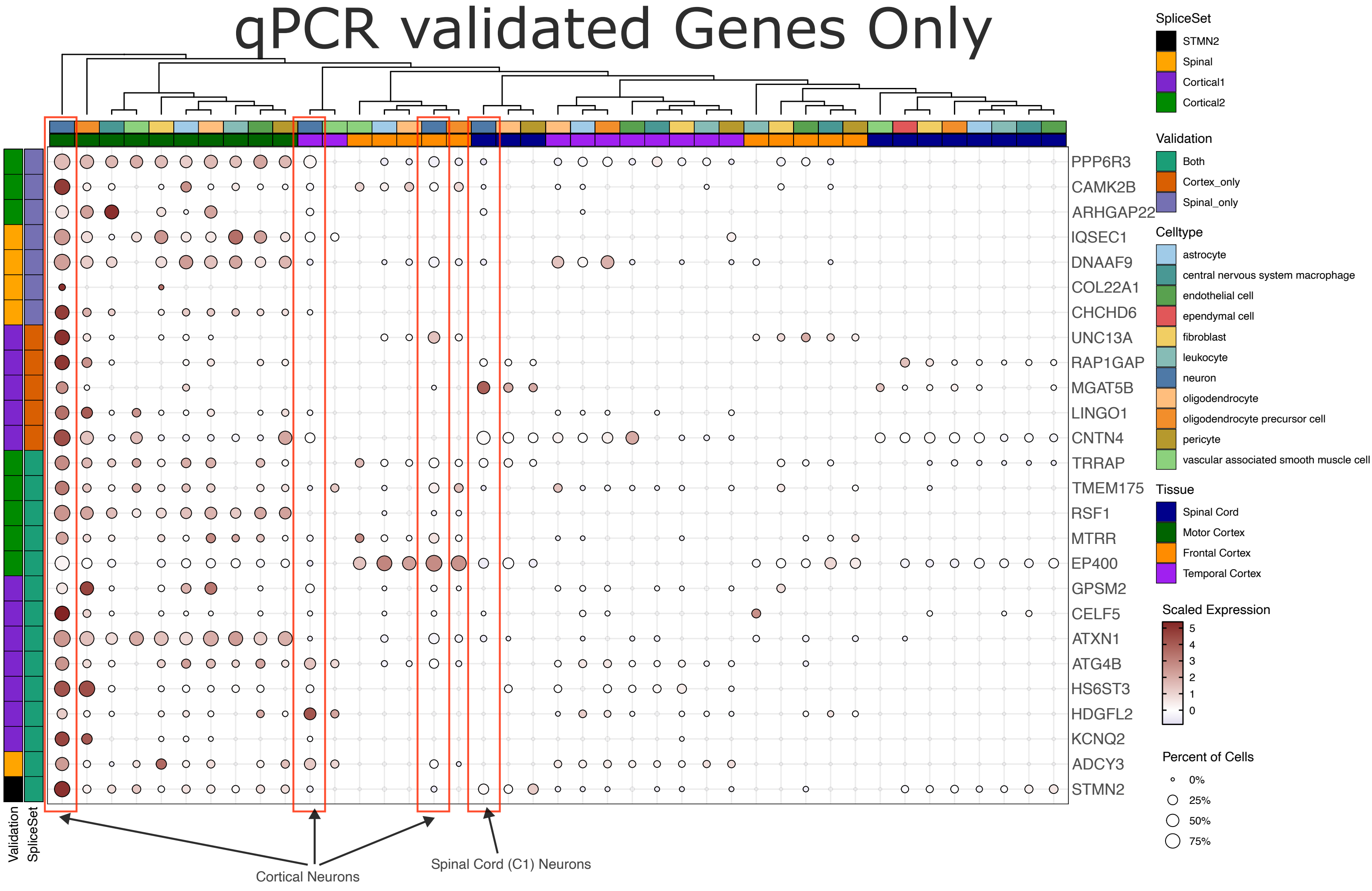


Online Resource 8. Expression of cryptic splice-harbouring genes across Spinal Cord and Frontal, Motor, and Temporal cortex in the Human Brain Cell Atlas. Dotplots representing intensity, and preponderance, of gene expression of genes harbouring cryptic splices. Dot size represents the percentage of cells, of each type and region, with colour indicating Z-normalized log2(size factor-normalized counts). Each row was Z-normalized independently. Size factors were first normalized within each anatomical region, then across regions, in order to account of systematic differences in library size across regions. Only genes harbouring cryptic splices which show significant elevation by qPCR are shown. Note the overall trend highlighted in Figure 3, that the cortical1 (and less so, cortical2) splice sets are broadly explained by greater expression in cortical regions (largely the motor cortex in this dataset) than spinal cord. Conversely the spinal splice set is not explained by differences in expression.

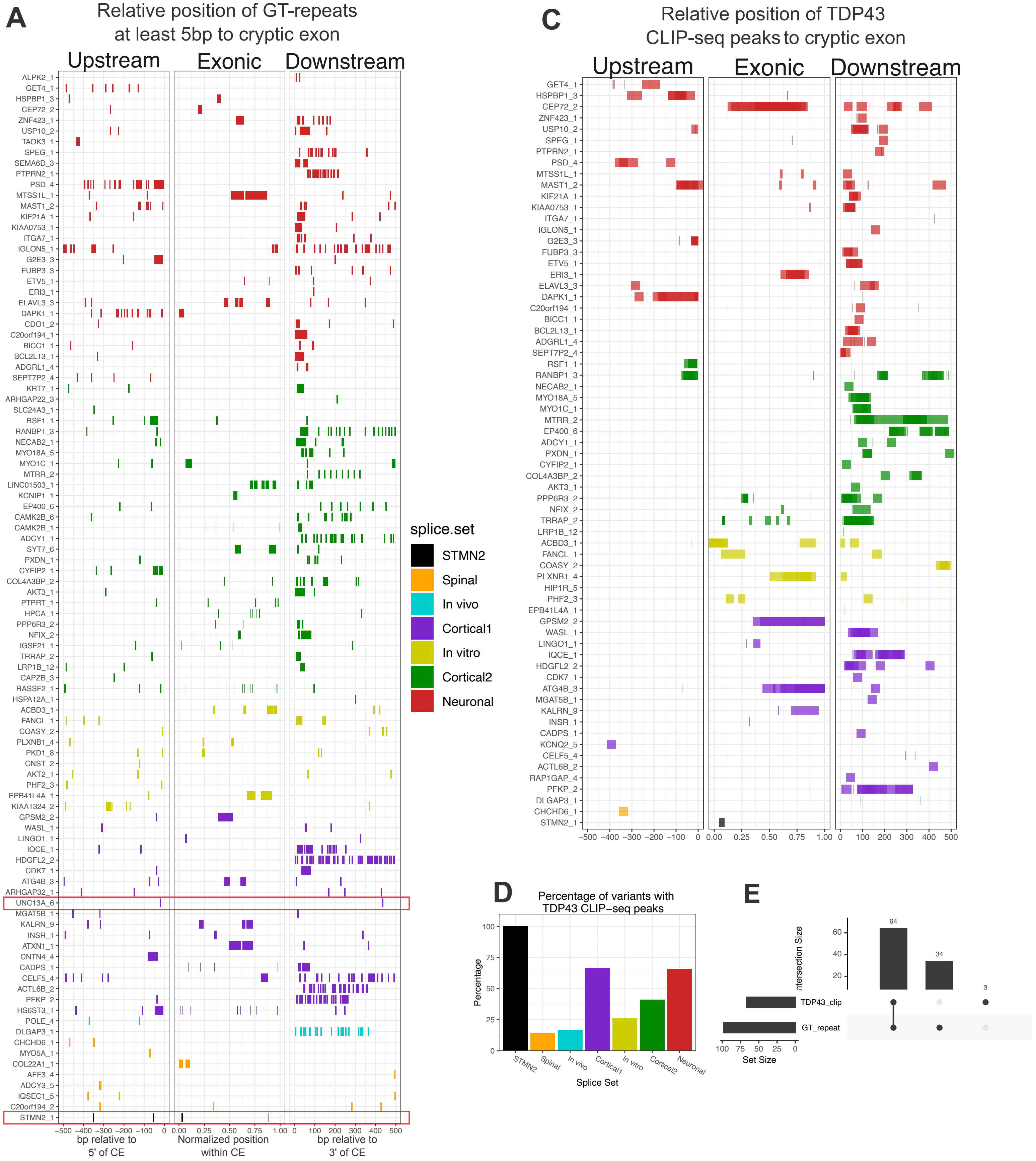


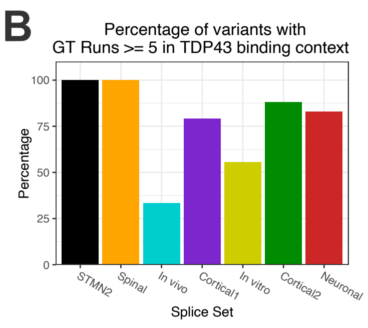


Online Resource 9. GT-repeats and TDP43 CLIP-seq peaks relative to cryptic exons of all cryptic splices. Positions of GT-repeats and CLIP-seq peaks were calculated relative to the boundaries of each cryptic exon, within a ±500bp window. Upstream (i.e. intronic space to the 5' end of the cryptic exon) positions were negative, intra-exonic positions were normalized on a 0-1 scale, representing the start and end of the exon, and downstream positions were positive. Positions were calculated in a strand-aware manner (i.e. upstream is always 5' of the cryptic exon). A - Relative positions of GT-runs with a length of 5bp or more. Note, the relatively short GT-repeats proximal to the UNC13A and STMN2 cryptic exons (red boxes). GT-run positions were coloured by splice set. Cryptic splices with no proximal >=5bp GT-runs were not visualized. B - Percentage of splice variants within each splice set containing a GT-run of at least 5 in the TDP43 binding context (area shown). C - Relative positions of TDP43 CLIP-seq peaks. D - Percentage of splice variants within each splice set containing a TDP43 CLIP-seq peak. Note the drop in spinal splice set percentage relative to the GT-runs in B. E. Overlap of cryptic splice variants with proximal TDP43 CLIP-seq peaks and/or GT-runs. Note, all CLIP-positive variants have GT-repeats, but not vice versa, highlighting a limited utility of GT-runs for inferring TDP43-dependence.

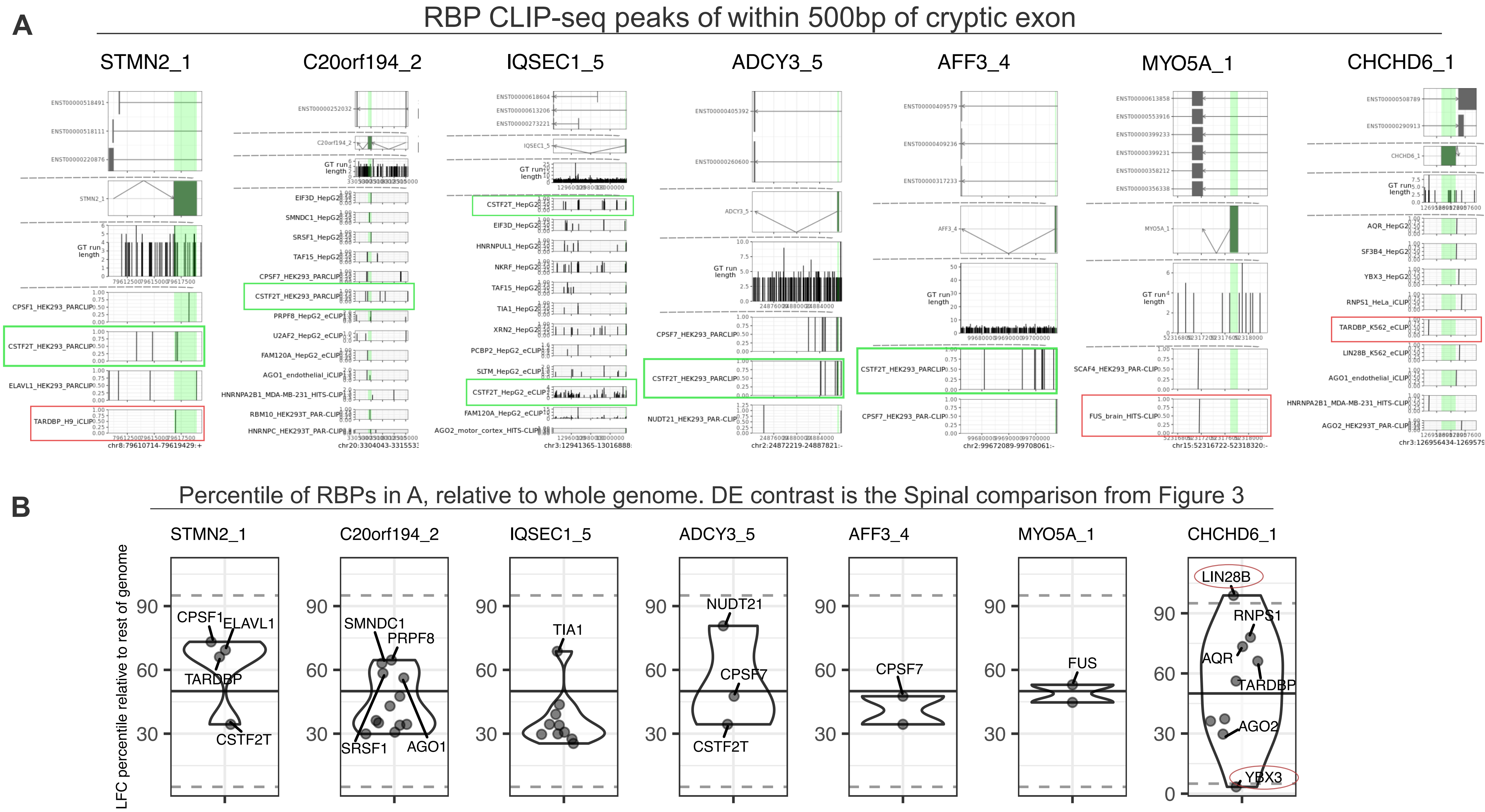


Online Resource 10. Exploring trans-regulatory RBP factors that may explain tissue-specific enrichment of the spinal splices. RNA binding protein (RBP) CLIP-seq peaks were collected from ENCODE, iONMF, and Muktherjee et al, 2019, to create a reference database of 570 CLIP-seq peaksets across 259 RBPs. Diverse CLIP modalities were captured, including CLIP, eCLIP, iCLIP, PARCLIP, and HITS-CLIP. RBPs with peaks proximal (500bp upstream/ downstream, based on Tollervey et al, 2011) to cryptic splice sites. To account for differences in coverage and intensity across experiments, peak signal was normalized on a 0-1 scale, representing the minimum and maximum signal of all peaks for that experiment. A - Relative position of CLIP-seq peaks from any RBP showing proximal peaks to the STMN2 cryptic splice, or the spinal splices. Each plot contains three sets of tracks, an upper track of canonical annotated transcripts, a second track showing the cryptic exon and associated splice junction(s), a third track showing the location of GT-rich repeats (strings of 4bp or longer) on the relevant genomic strand, and a lower bank of tracks for all RBP CLIPseq experiments with proximal peaks. Cryptic exon coordinates are highlighted in green. Note the TARDBP peak in the cryptic exon of STMN2, and upstream of CHCHD6, as well as a FUS peak in MYO5A (red boxes). Green boxes highlight CLIP peaks for CSTF2T, which are proximal to the STMN2, ADCY3, IQSEC1, C20orf194, and AFF3 cryptic exons. B - Relative expression percentiles of RBPs with proximal peaks to cryptic exons for each cryptic splice variant in A. Percentiles were calculated as in Figure 3, with the “Spinal” comparison being shown: Controls from cervical, thoracic, and lumbar spinal cord versus frontal, temporal, motor cortex, RiMOD. Liu (sorted TDP43+ nuclei), and all in vitro scrambled controls. STMN2-proximal RBPs showed no clear enrichment (>95th or <5th percentile) in spinal cord, as expected given its context-independence. Other spinal splices also showed little enrichment, with the exception of CHCHD6 which had proximal peaks from LIN28B (elevated in spinal cord) and YBX3 (decreased in spinal cord).

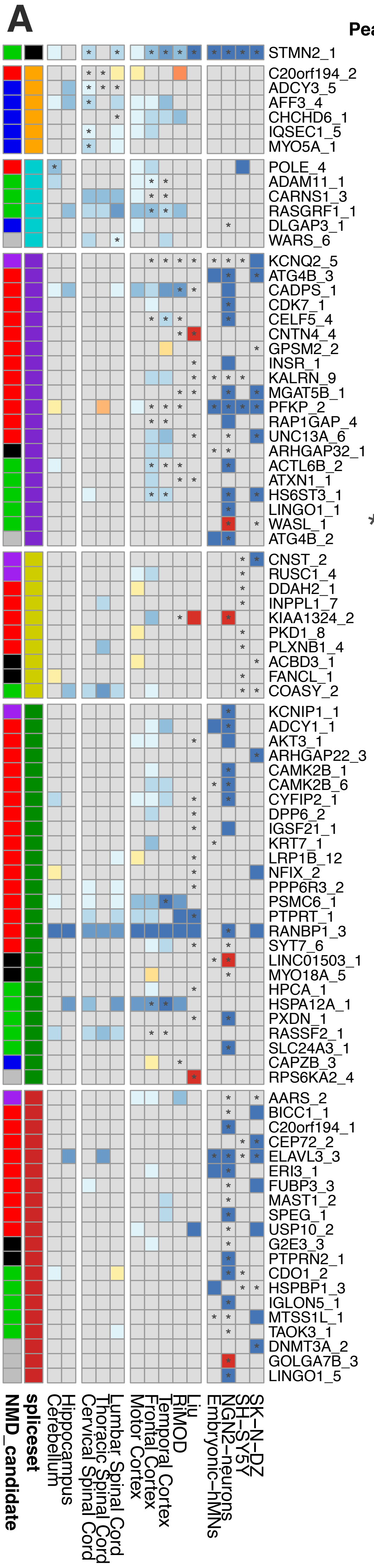


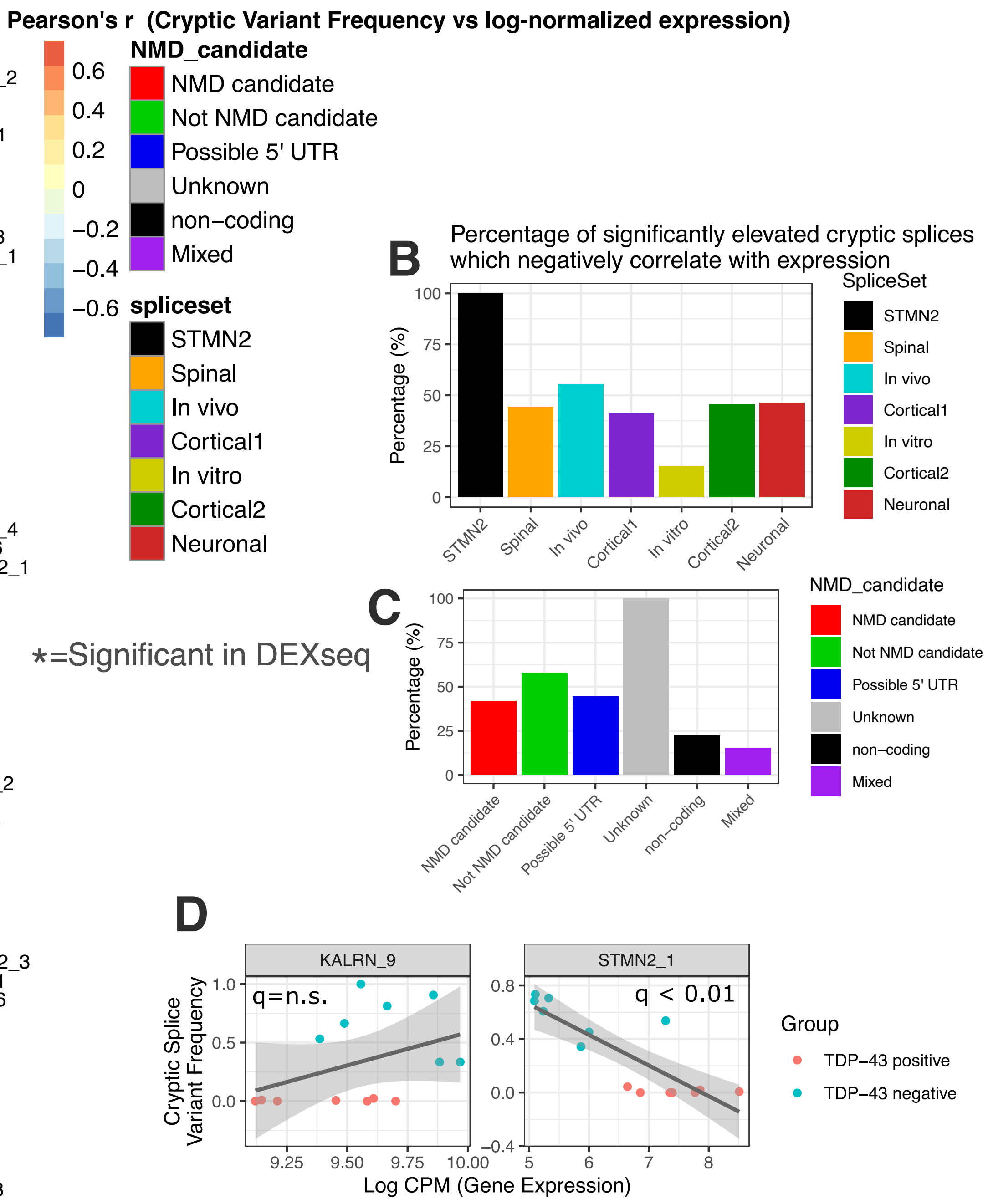


Online Resource 11. Relationships between cryptic splice inclusion and gene expression. A - Heatmap showing significant (5% FDR) Pearson correlations between cryptic splice variant frequency and gene expression. Contexts where significant DEXseq elevations in variant frequency (i.e. where cryptic splices were elevated in the initial meta-analysis), and indicated by asterisks. B - Barplot showing the percentage of significant DEXseq hits which show significant negative correlations between variant frequency and gene expression (e.g. for STMN2, 100% of contexts with DEXseq-significant elevations in cryptic splicing also show reductions in gene expression), aggregated by splice set. Note: for most splice sets this percentage is less than or equal to 50%, suggesting that many cryptic splices do not result in the downregulation of their respective gene. C - Same data as in B, but aggregated by predicted splice effect. D - Scatterplots showing examples of the correlations underlying A. Both plots show data from Liu et al (sorted TDP43+/- nuclei), highlighting a a n.s. correlation between cryptic variant frequency and expression for KALRN (and indeed a slight positive tendency in the correlation), and a negative correlation for STMN2 (consistent with a wide array of literature reports).

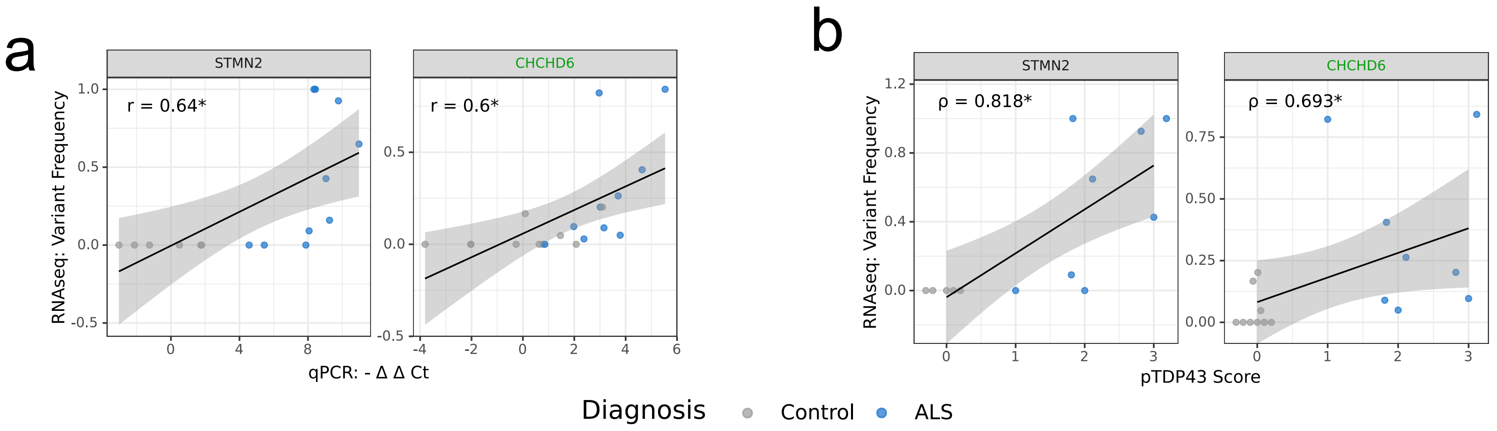


Online Resource 12. Detection of STMN2 and CHCHD6 cryptic splices in a novel spinal cord RNAseq cohort. A - pTDP43 IHC scores (semi-quantitative) for control and ALS spinal cord sections. Note that 2 ALS and 1 control section had uninterpretable IHC (thus are NA values in this and subsequent plots). B - Correlation of qPCR and RNAseq quantifications of STMN2 and CHCHD6 cryptic splices (i.e. those which were detected in both qPCR and RNAseq). Significant Pearson correlation coefficients are labeled on each plot. *p< 0.05.

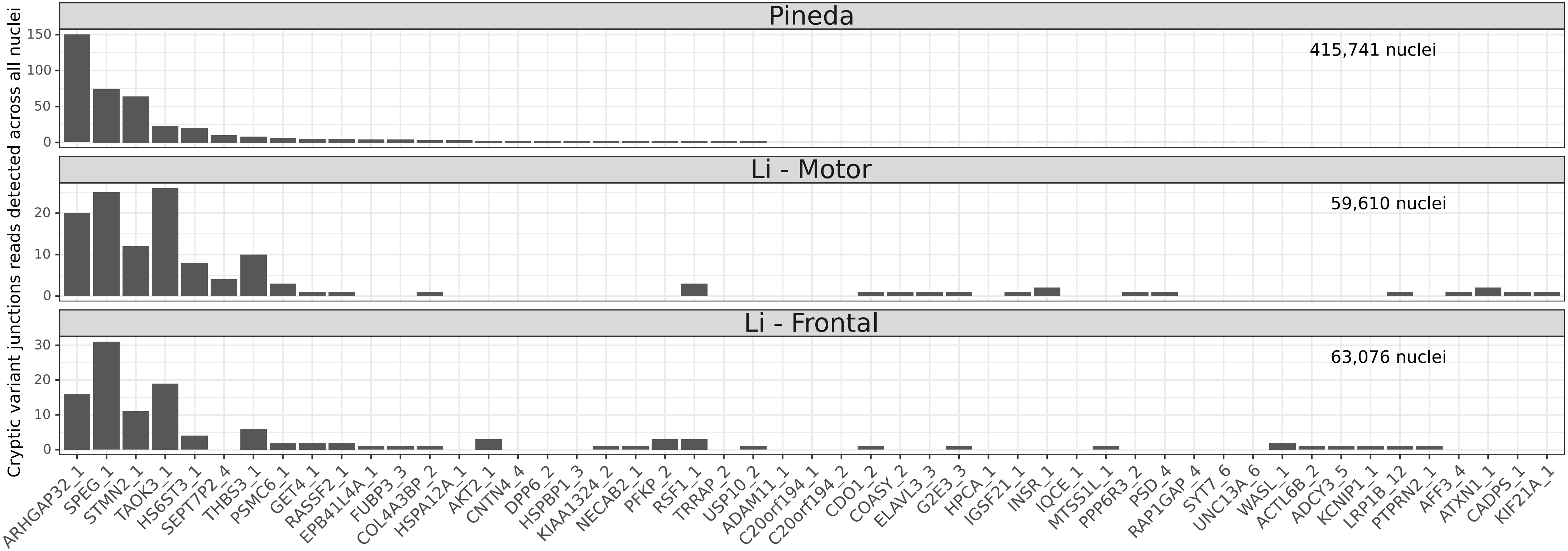


Online Resource 13. Distribution of cryptic splice junction counts across all cells snRNAseq datasets. Histogram showing the total number of cryptic splice junctions detected across all cells in Pineda (upper), Li motor cortex (middle), and Li frontal cortex (lower). Total nuclei included in each dataset are printed on each plot. Note the greater total number of cryptic splice junctions in Pineda, and the ~7-fold greater number of nuclei.

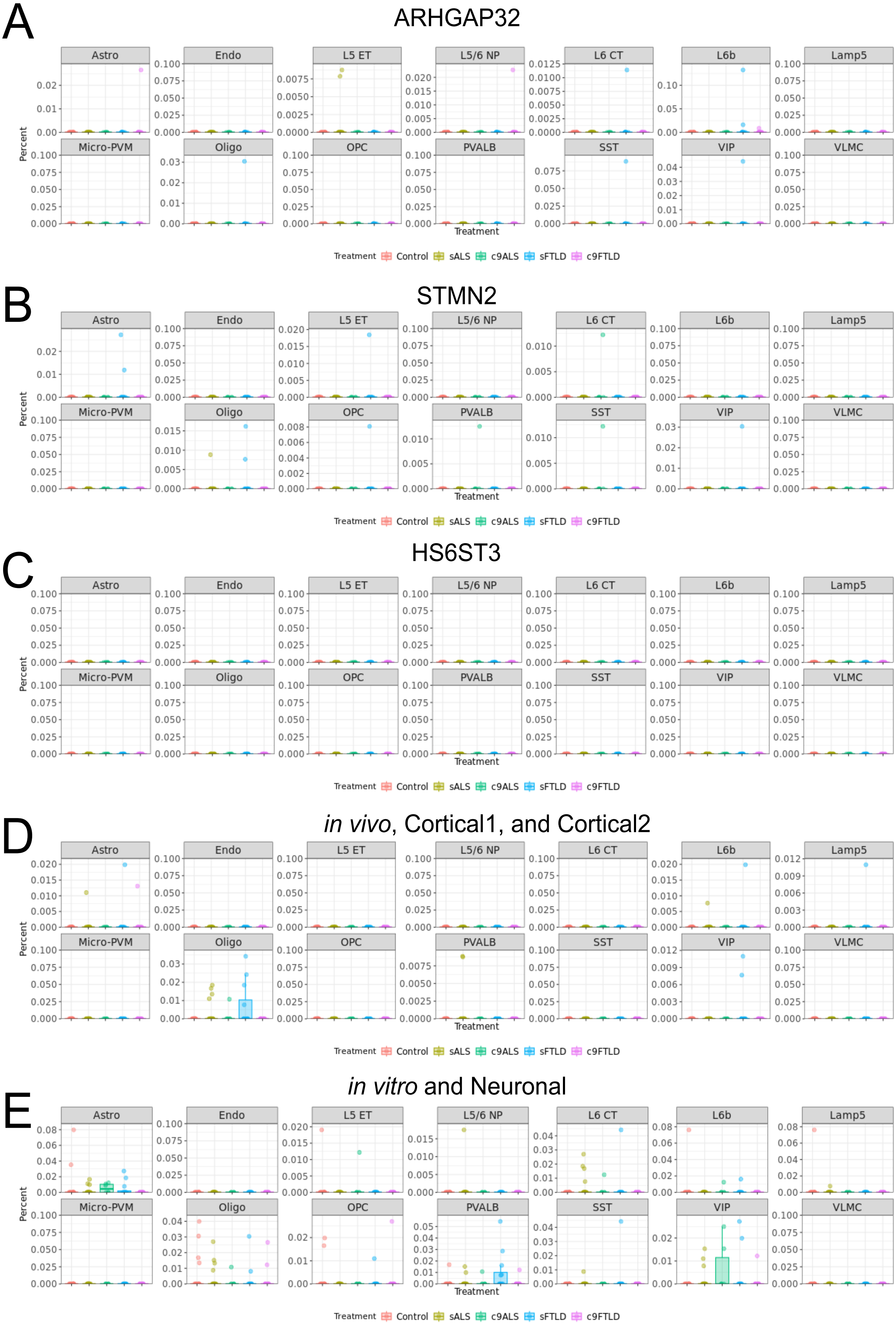


Online Resource 14. Cryptic cellularity plots of all cell-types identified in analysis of Pineda et al dataset. All plots show cryptic cellularity for each cell-type (i.e. the percentage, per sample, of cryptic splice-positive nuclei in each cell-type population), y-axis shows values >1%, not fractions. A - ARHGAP32 cryptic cellularity. B - STMN2 cryptic splice. C - HS6ST3 cryptic splice. D - Cryptic cellularity of the remaining detected in vivo, cortical1, and cortical2 cryptic splice variants. E - Cryptic cellularity of the remaining detected in vitro and neuronal (i.e. in vitro and Liu-enrichment) cryptic splice variants.

### Li *et al*, Motor Cortex

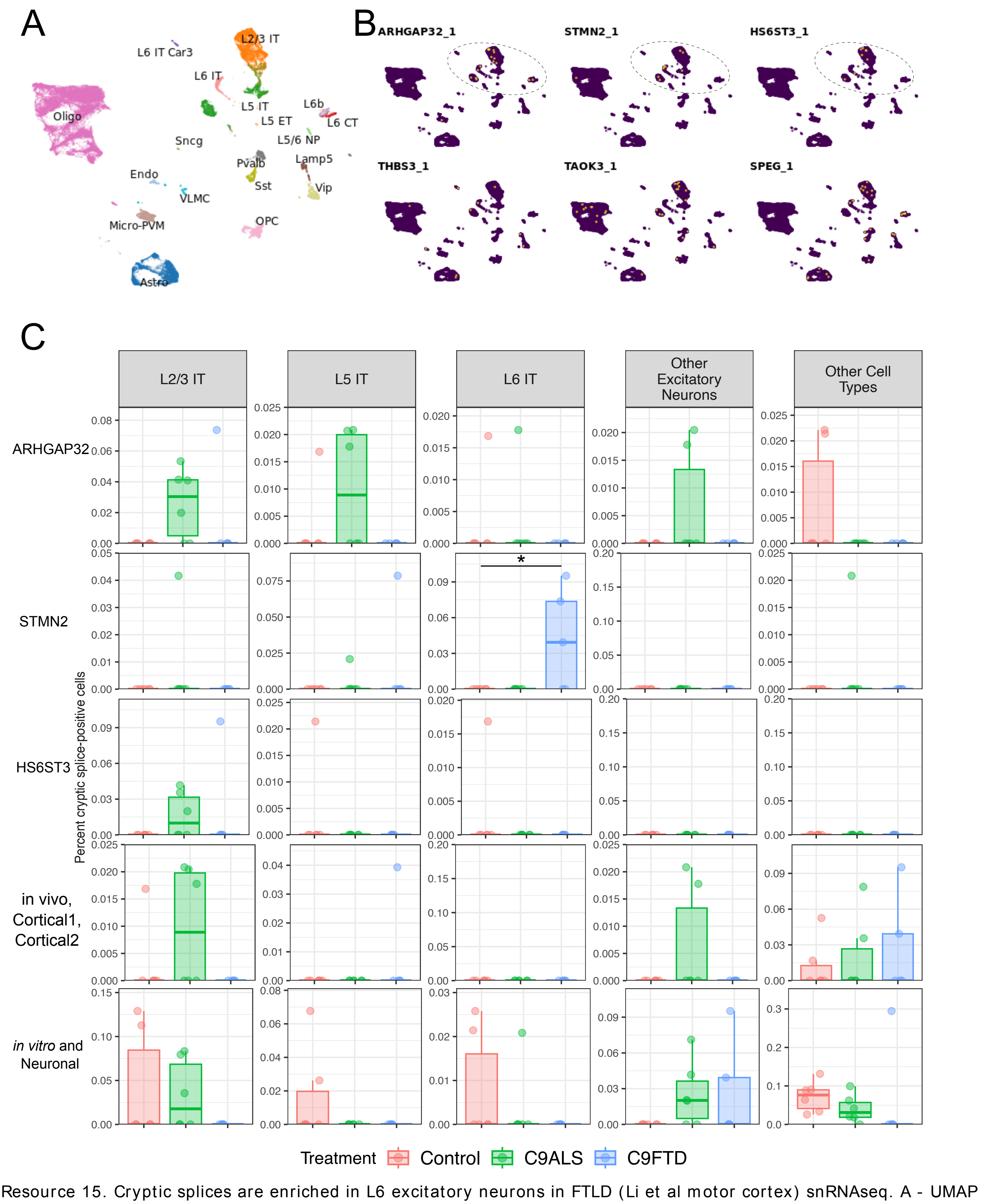


Online Resource 15. Cryptic splices are enriched in L6 excitatory neurons in FTLD (Li et al motor cortex) snRNAseq. A - UMAP embedding showing identified cell populations in analysis of snRNAseq data from Li et al (Motor Cortex). B - UMAPs showing cryptic splice-harboring cells in yellow. Dashed circle highlights populations of excitatory neurons harboring the majority of cryptic splicing. C - Cellularity plots showing the percentage of cryptic-positive cells, for each cell-type, relative to the total number of detected cells (per donor), y-axis shows values > 1%, not fractions. "Other excitatory" combined L5 ET, L5/6 NP, L6b, and L6 CT populations, Interneurons combined Lamp5, VIP, SST, and PVALB populations, with all other populations combined in "Other Cell Types". Population-wise cryptic cellularity for "Other Cell Types" is visualized in Online Resource 16. Each dot represents an individual donor. Significance was derived from differential abundance testing using edgeR, to account for potential composition effects. * q< 0.05, ** q< 0.01, *** q< 0.001.

### Li *et al*, Motor Cortex

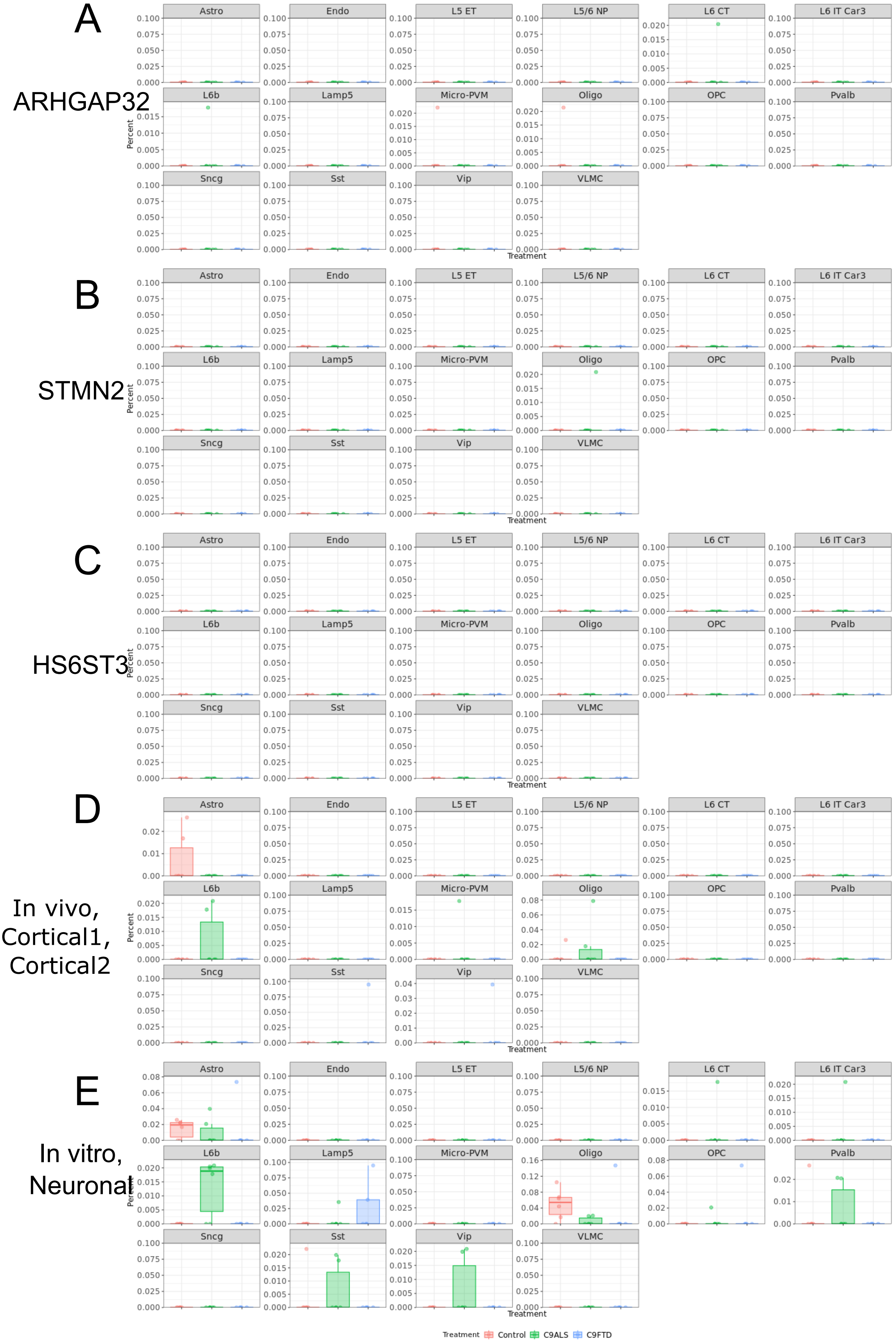


Online Resource 16. Cryptic cellularity plots of all cell-types identified in analysis of Li et al motor cortex. All plots show cryptic cellularity for each cell-type (i.e. the percentage, per sample, of cryptic splice-positive nuclei in each cell-type population), y-axis shows values >1%, not fractions. A - ARHGAP32 cryptic cellularity. B - STMN2 cryptic splice. C - HS6ST3 cryptic splice. D - Cryptic cellularity of the remaining detected in vivo, cortical1, and cortical2 cryptic splice variants. E - Cryptic cellularity of the remaining detected in vitro and neuronal (i.e. in vitro and Liu-enrichment) cryptic splice variants.

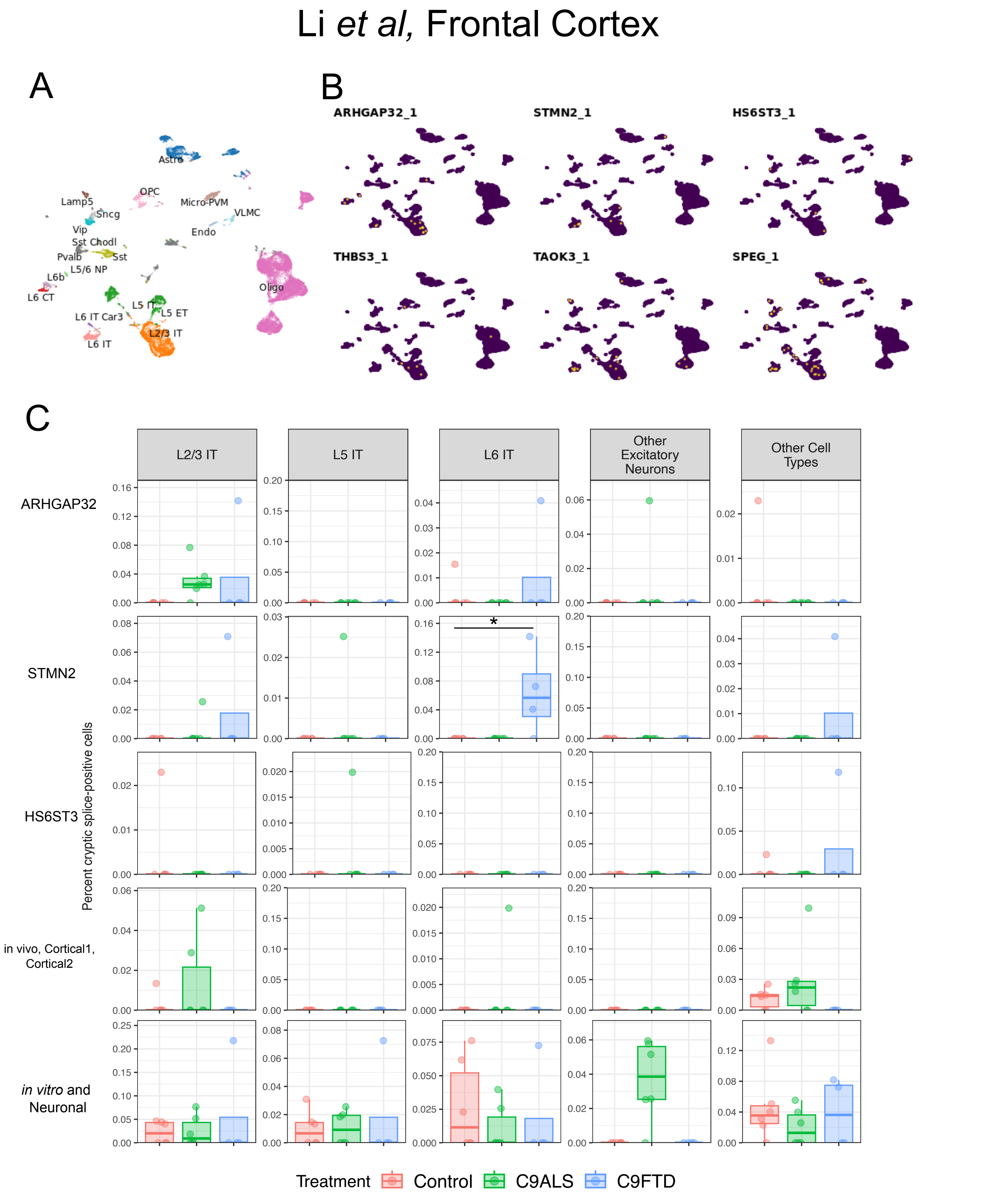


Online Resource 17. Cryptic splices are enriched in L6 excitatory neurons in FTLD (Li et al frontal cortex) snRNAseq. A - UMAP embedding showing identified cell populations in analysis of snRNAseq data from Li et al (Frontal Cortex). B - UMAPs showing cryptic splice-harboring cells in yellow. Dashed circle highlights populations of excitatory neurons harboring the majority of cryptic splicing. C - Cellularity plots showing the percentage of cryptic-positive cells, for each cell-type, relative to the total number of detected cells (per donor), y-axis shows values > 1%, not fractions. “Other excitatory” combined L5 ET, L5/6 NP, L6b, and L6 CT populations, Interneurons combined Lamp5, VIP, SST, and PVALB populations, with all other populations combined in “Other Cell Types''. Population-wise cryptic cellularity for “Other Cell Types'' is visualized in Online Resource 18. Each dot represents an individual donor. Significance was derived from differential abundance testing using edgeR, to account for potential composition effects. * q< 0.05, ** q< 0.01, *** q< 0.001.

### Li *et al*, Frontal Cortex

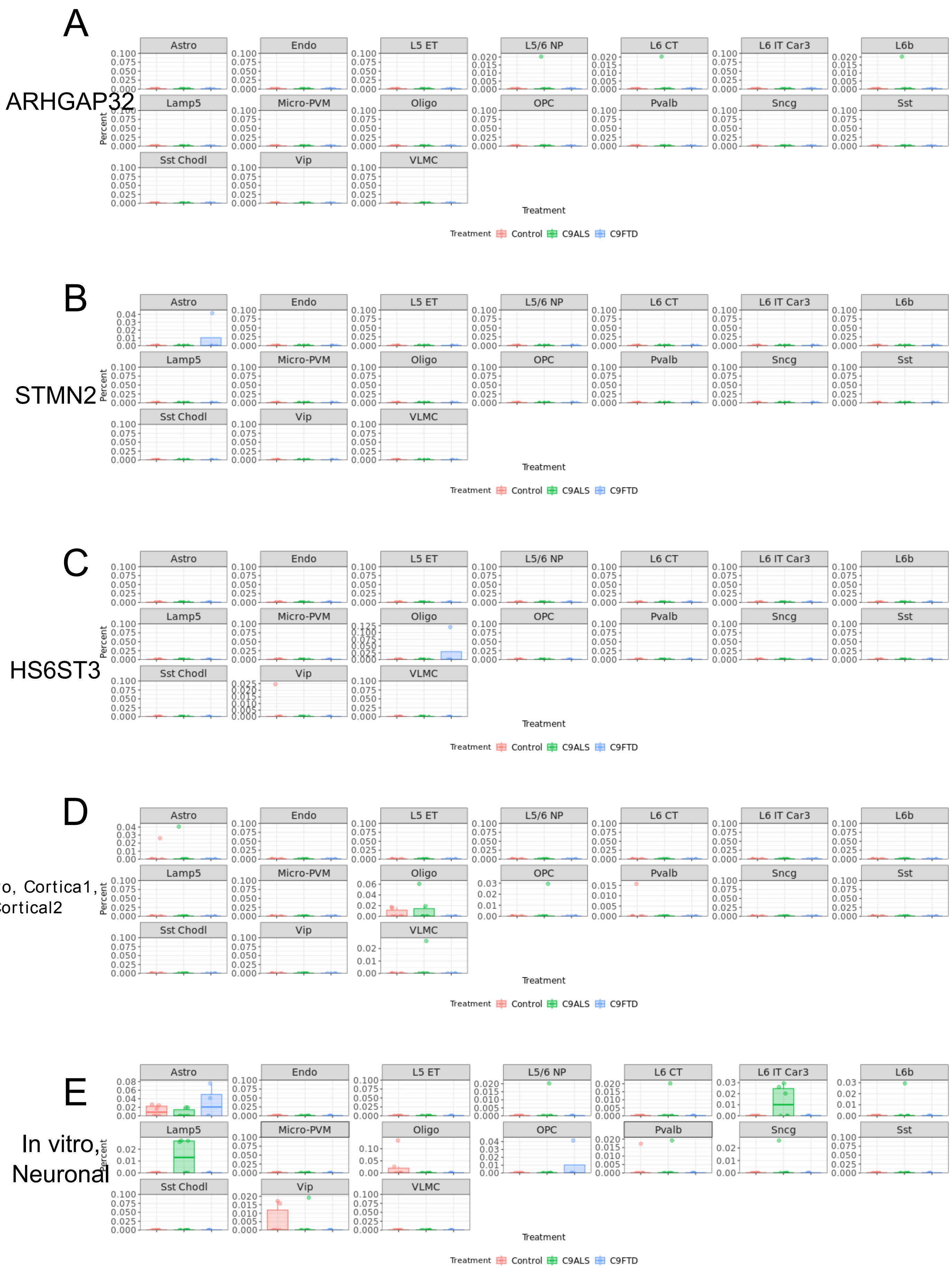


Online Resource 18. Cryptic cellularity plots of all cell-types identified in analysis of Li et al frontal cortex. All plots show cryptic cellularity for each cell-type (i.e. the percentage, per sample, of cryptic splice-positive nuclei in each cell-type population), y-axis shows values > 1%, not fractions. A - ARHGAP32 cryptic cellularity. B - STMN2 cryptic splice. C - HS6ST3 cryptic splice. D - Cryptic cellularity of the remaining detected in vivo, cortical1, and cortical2 cryptic splice variants. E - Cryptic cellularity of the remaining detected in vitro and neuronal (i.e. in vitro and Liu-enrichment) cryptic splice variants.

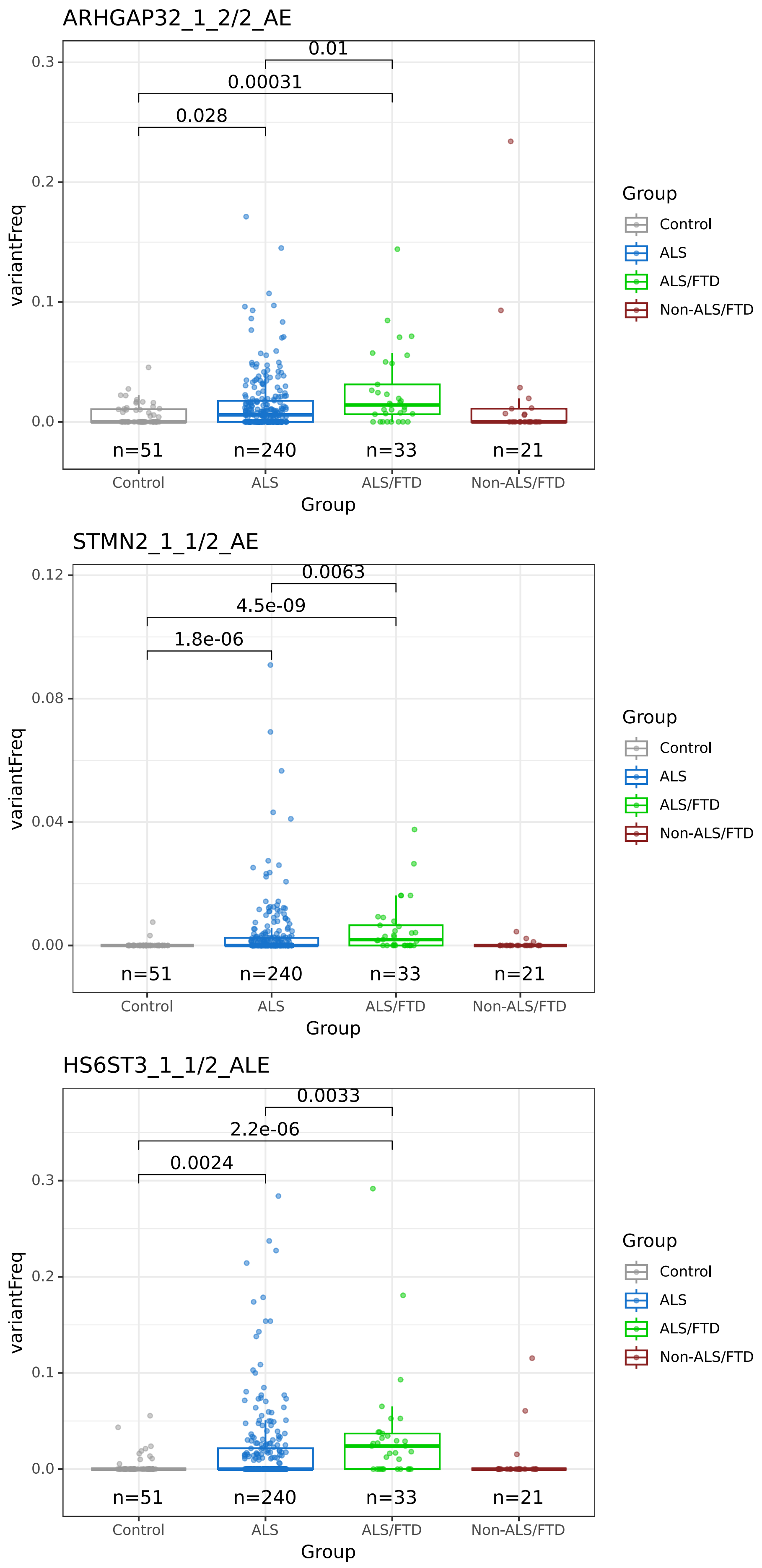


Online Resource 19. Variant frequency plots of ARHGARP32, STMN2, and HS6ST3 cryptic splices in motor cortex bulk RNAseq. Plots showing variant frequency across diagnostic groups, for the motor cortex only. Unpaired t-test p-values shown. Note that across all three cryptic splices elevated in snRNAseq data, both ALS and ALS/FTD groups are elevated above controls, though ALS/FTD are further increased above ALS alone.
